# The signaling mechanism of Hcy-induced atrial fibrosis mediated by TRPC3

**DOI:** 10.1101/583740

**Authors:** Lu Han, Yanhua Tang, Yanqing Wu, Xiaoshu Chen, Kui Hong, Juxiang Li

**Author notes:** **Corresponding author: Juxiang Li**, MD & PhD; **Yanhua Tang**, MD & PhD. **Postal Address:** Department of Cardiovascular Medicine, the Second Affiliated Hospital of Nanchang University, Minde Road No.1, Nanchang of Jiangxi, 330006, China.

## Abstract

**Background:** High plasma levels of homocysteine (Hcy) are regarded as a risk factor for atrial fibrillation (AF), which is closely associated with the pathological consequence of atrial fibrosis and can lead to heart failure with a high mortality rate; Currently, there is no effective therapy for preventing atrial fibrosis, owing to a lack in fundamental understanding of the underlying mechanism. Here, we show that atrial fibrosis is mediated by the relationship between canonical transient receptor potential 3 (TRPC3) channels and sirtuin type 1 (SIRT1) under the stimulation of Hcy.

**Methods:** The left atrial appendage was obtained from patients with either sinus rhythm (SR) or AF, who underwent cardiothoracic surgery, and used to evaluate the relationship between the concentration of Hcy and a potential mechanism of cardiac fibrosis mediated by TRPC3 and SIRT1. We next performed transverse aortic constriction (TAC) in mouse to investigate the relationship. The mechanisms underlying atrial fibrosis involving TRPC3 and SIRT1 proteins were explored by co-immunoprecipitation (co-IP), bio-layer interferometry (BLI) and lentivirus transfection experiments. Quantitative polymerase chain reaction (qPCR) and western blotting (WB) were performed to analyse gene and protein expression, respectively.

**Results:** The majority of AF patients displayed atrial fibrosis, as demonstrated by Masson staining and immunohistochemistry. In the mouse model of TAC, more severe fibrosis was detected in the high-Hcy diet (HH) group, compared to NH mice; and the duration of induced AF was longer in the HH groups than in the normal diet (NH) group. Moreover, the HH group exhibited higher expression levels of TRPC3 and related fibrosis proteins, such as TGF-ß and Col-I, than the NH group, despite also showing a higher level of SIRT1 was observed. The activator of SIRT1 (Resveratrol, Res) attenuated the enhancement of TRPC3 and decrease in SIRT1 observed in the HH group. Further cell culture experiments confirmed that Hcy could promote the proliferation and differentiation of fibroblasts, the up-regulation of TRPC3, and the decrease in the protein level of SIRT1. Ultimately, the results of Co-IP and BLI indicated a direct interaction between TRPC3-C terminal domain (569-863) and SIRT1 proteins, in which the two proteins are antagonistic and in combination regulate the pathogenesis of atrial fibrosis.

**Conclusions:** The higher level of atrial fibrosis were observed in the HH mouse group, compared with the NH mice group, Such results suggest that AF patients may be more susceptible to atrial fibrosis and possess a high probability of progressing to hyperhomocysteinemia. Moreover, our findings are consistent with the hypothesis that TRPC3 channel up-regulation leads to abnormal accumulation of collagen, with the down-regulation of SIRT1 as an aetiological factor of high Hcy, which in turn predisposes to atrial fibrosis and strongly enhances the possibility of AF.

## 1. Introduction

Atrial fibrosis is an inevitable pathological mechanism in several different cardiac diseases, especially AF, which can lead to cerebral apoplexy and frequent hospitalizations ^[2,3]^. We have previously demonstrated that homocysteine (Hcy) is significantly associated with the recurrence of AF after radiofrequency ablation ^[1]^. Elevated levels of Hcy in plasma are an independent risk factor for cardiovascular disease, and most cardiovascular conditions accompanied by hyperhomocysteinemia is significantly correlated with cardiac fibrosis. This indicates that high Hcy may promote atrial fibrosis ^[4]^. Structural remodelling in atrium can be easily detected in paroxysmal and permanent AF rather than pathological changes in ventricle. Similarly, cardiac fibrosis is accompanied by HF due to the accumulation of collagen fibres, such that patients with AF have HF with a preserved ejection fraction (HFpEF) rather than HF with reduced ejection fraction (HFrEF), accounting for 30% of cases ^[5–7]^. These findings suggest that the atrial fibrosis is associated with collagen metabolism and is involved in the mechanitsm of HF. Furthermore, hyperhomocysteinemia is a pathological feature in the aetiology of atrial fibrosis, although the underlying mechanisms remain unclear ^[8]^. Canonical transient receptor potential receptor 3 (TRPC3) is an indispensable factor in regulating the mechanisms of fibrosis development, and in promoting the transition of fibroblasts into myofibroblasts with an adverse influence on the modulation of collagen ^[9]^. On the other hand, sirtuin-1 (SIRT1) appears to function as an anti-fibrosis protein. Recently, the progression of myocardial fibrosis has been shown to simultaneously activate the renin-angiotensin system (RAS). This causes myocardial apoptosis through the TGF-ß pathway, and controls the aggregation of monocytes and fibroblasts, leading to the down-regulation of the mitochondrial nicotinamide adenine dinucleotide dependent deacetylase called SIRT1 ^[10]^. Whether TRPC3 and SIRT1 can control and modulate the fibrosis system to reciprocally affect cardiac structural remodelling remains unclear. In an attempt to address this issue, we hypothesize that TRPC3 is a novel regulator of SIRT1 in modulating the TGF-ß pathway. Although it remains unknown how TRPC3 regulates TGF-ß signalling, our results suggest that TRPC3 could indirectly or directly affect the activation of SIRT1, which subsequently regulates the TGF-ß signalling pathway ^[11]^. Therefore, our study aimed to elucidate whether SIRT1 is directly involved in the above signalling pathway and its role in fibroblast proliferation and differentiation under high Hcy conditions.

## 2. METHOD DETAILS

### 2.1 Human tissue specimens

Tissue samples were obtained from the left atrial appendage of 30 patients. All samples were collected at the Second Affiliated Hospital of Nanchang University between January 2016 and September 2017. The sinus rhythm (SR) group comprised patients with sinus rhythm and preserved left ventricular function (n=12); the atrial fibrillation (AF) group comprised patients with atrial fibrillation and rheumatic mitral stenosis (n=18). The study protocol was approved by the Institutional Review Board of the Second Affiliated Hospital of Nanchang University (Permit Number: 2016-022). Detailed clinical and pathological information on the patients is summarized in Table1.

### 2.2 Animal model

All animal experiments were approved by the SLAC Labomouseory Animal Co. Ltd, Hunan, China. Mice were kept on a 12 h light / 12 h dark cycle at a room temperature of 20-25 degrees, with a relative humidity of 40-70%. Baseline information on male C57B6 mice (n=60) was detected by transthoracic echocardiography. All experienced mice underwent transverse aortic constriction (TAC) at four weeks of age following randomization. Mice in the high-Hcy (HH) diet group were fed a high-Hcy diet (AIN-76A + 4% methionine with irradiation, ReadyDietech, China) to induce hyperhomocysteinaemia with heart failure (HF) (n=25), and mice in the non-Hcy (NH) diet group were fed a normal diet (n=25). The majority of the mice in the NH and HH groups showed HF signs after 7 weeks of age. The HH+Res group included mice that underwent TAC with a HH diet that were subjected to a single intraperitoneal injection of 20mg/kg/day resveratrol (Res) for 21 days ^[12]^. The mice in the HH group received the same dose of saline by intraperitoneal injection and underwent sham surgery. These mice were fed a folic acid and high-Hcy mixed diet classed as HH+FC group. (Table 3)

### 2.3 Transthoracic echocardiography

Mice were anaesthetized with 5% isoflurane for transthoracic echocardiography which was performed using a Vevo2100 imaging system (VisualSonics, Toronto, Canada). Ejection fraction (EF) was regarded as a systolic parameter, and E/A and E/E’ ratios were regarded as diastolic markers via baseline echocardiography. Pulsed-wave Doppler and tissue Doppler were performed to detect the peak ratio of E/A and E/E’ in the three groups at 4,7,16 weeks. Left ventricular (LV) end-diastolic volume (EDV) and end-systolic volume (ESV) were obtained by the Simpson method of disks. Ejection fraction was calculated as EF (%) = (EDV–ESV)/EDV×100% and was used to determine systolic function from images in the parasternal short-axis view as previously described ^[13]^. Left ventricular end-diastolic diameter (LVEDD) and end-systolic diameter (LVESD) were recorded. Fractional shortening (FS) was evaluated with the following formula: FS (%) = (LVEDD–LVESD) /LVEDD×100%. The E/A ratio was determined to evaluate diastolic function in pulsed-wave Doppler mode ^[13,14]^. TAC mice demonstrated diastolic dysfunction by echocardiography, including both the NH and HH groups.

### 2.4 Surface electrocardiography

Surface electrocardiography (ECG) was recorded prior to *in vivo* arrhythmia induction studies for HH, NH and SH mice at 16 weeks of age (SLAC Labomouseory Animal Co. Ltd, Hunan, China) ^[15]^. The PR interval, QRS dimension, QT interval, and RR interval were measured three times and averaged on MedLab6 software (Biological signal acquisition and processing system, Beijing, China).

### 2.5 Histopathological examination

Atrial tissues were fixed in 4% paraformaldehyde, embedded in paraffin wax and underwent different staining methods, including haematoxylin-eosin (H+E), Masson and immunohistochemical staining, each according to the manufacturer’s instructions (Sigma-Aldrich). Primary antibodies used were anti-TGF-ß (5 μg/ml) (CST, 3711) and anti-collagen-I (5 μg/ml) (Thermo Fisher Scientific, PA5-16697). For electron microscopy, heart tissues were fixed with special fixative at 4°C for 2-4 h and rinsed 3 times with 0.1 M phosphate buffer (PB). Next, the tissues were fixed with 1% osmium acid, rinsed with 0.1 M PB (pH=7.4), and dehydrated with ascending ethanol concentrations for 15 min each time. The tissues were immersed in Epon812 resin/acetone (1:1) for 2-4 h, immersed in fresh Epon 812 resin for 30 min, and embedded for convergence overnight at 60°C. The sample was cut to an approximately 60-80 nm thickness using a ultrathin section machine, subjected to uranium-lead double-staining (2% uranium acetate saturated alcohol solution, lead citrate, 15 min staining), and observed on a transmission electron microscope (HT-7700, Hitachi, Tokyo, Japan).

### 2.6 Immunofluorescence and immunohistochemistry

For conventional fixation, cells were immersed in 4% polyoxymethylene and 0.1% Triton X-100, washed 3 times with sterile phosphate-buffered saline (PBS), and incubated with primary antibodies in PBS containing 5% bovine serum albumin (BSA) at 4°C overnight. After the samples were washed 3 times with PBS, they were incubated with Alexa488- or 564-conjugated goat anti-rabbit/mouse secondary antibodies at room temperature for 1.5 h. Finally, each group of cells was stained with 4′,6-diamidino-2-phenylindole (DAPI). To detect the localization of SIRT1 and TRPC3 under Hcy stimulation, HA-SIRT1 and Flag-TRPC3 were overexpressed in HEK 293 cells, which were treated with or without Hcy (500 μmol/L) for 48h. Representative images were acquired using a BWS435 confocal microscope and LAS AF Lite software was used for professional image analysis (Zeiss, Germany).

Tissue sections were incubated in 10 mM citric acid buffer (pH 6.0) at boiling temperatures for 8 min, placed in PBS (pH7.4), washed 3 times on a decoloring shaker, and incubated with 3% BSA for 30 min at room temperature. Sections were incubated with primary antibody (1:100) at 4°C in a wet box overnight, and the antigen-antibody reaction was observed using a horseradish peroxidase (HRP)-conjugated secondary antibody after incubation for 50 min. Image acquisition was performed using an XSP-C204 microscope (CIC).

### 2.7 Cell lines and treatment

MCF and HEK-293T cell lines were purchased from Procell Life Science and Technology (CP-M074 MCF) and American Type Culture Collection (CRL-3216 293T) between 2016 and 2017. The cells used in this study were cultured in DMEM medium (#11960051) (Gibco, CA, USA) supplemented with 10% foetal bovine serum (#10099-141) (Gibco, CA, USA). Pyrazole-10 (HY-19408), Resveratrol (HY-16561) and Salermide (HY-101073) were purchased from MedChemExpress (USA), and Hcy (H-4628) was obtained from Sigma-Aldrich (St. Louis, MO).

### 2.8 Isolation of mouse atrial fibroblast

Mouse atrial fibroblasts were collected and cultured from neonatal C57B6 mice at 0-2 days (Slake, Hunan, CHINA), weighing 3-5 g, in order to identify the relationship between atrial fibrosis and Hcy. The specific methods for mouse atrial fibroblast culturing are as follows. Firstly, mice were disinfected and the neonatal heart was rapidly removed. Left and right ventricles with a magnifying glass, and ventricular tissue was removed. The myocardial tissue was digested by addition of a trypsin/Collagenase II mixture 3 times, followed by treatment with a serum-containing medium that inhibited enzyme activity. The cell suspension was centrifuged (500 g, 5 min) it, collected and plated. Cells were allowed to adherence for 2 hours in a CO_2_ incubator.

### 2.9 Lentivirus infection

The Lenti-TRPC3-shRNA Tet-On construct (VB171023-1015pst) for TRPC3 knockdown was generated, packed, and purified by VectorBulider (Shanghai, China). The TRPC3-shRNA target sequence was 5’-CCTAAGGTTAAGTCGCCCTCG-3’. The final product was Puro-U6>mTrpc3. The empty vector used was puro-U6>Scramble. The scramble target sequence was 5’-CCTAAGGTTAAGTCGCCCTCG-3’. For SIRT1 overexpression, the promoter of EF1A was inserted into a lentiviral vector, and the ORF (mSIRT1 (NM_019812.3)) was subcloned into VB171024-1030vur. An adenovirus vector containing a GFP reporter was transfected into cells for efficient selection of infected cell lines. For lentivirus infection, fibroblasts were plated in T25 culture flasks at 4 × 10^3^ cells/cm^2^ and infected with Ad-TRPC3-shRNA, Ad-SIRT1-overexpression or empty vector (Ad-GFP) at a multiplicity of infection of 20 (MOI=20). The infection efficiency was determined by GFP fluorescence intensity (90-95%).

For the animal experiments, male C57BL/6 mice were injected with 100 μl purified lentivirus at one week of age via the caudal vein (SLAC Labomouseory Animal Co. Ltd, Hunan, China), with either Ad-GFP, Ad-SIRT1-overexpression or lentivirus plasmids inhibiting TRPC3 (Ad-TRPC3-shRNA), 12 h before TAC or sham surgery ^[16]^. Mice in the HH+vector or NH+vector group underwent TAC surgery and lentivirus injection with a high-Hcy or normal diet, respectively. Mice were administered 2 μg/ml doxycycline and 5% sucrose in sterile drinking water ^[17]^. After 3 weeks, heart tissues were obtained for IHC and WB analysis. All animal experiments were approved by the Ethic Committee of the Second Affiliated Hospital of Nanchang University (Permit Number: 2016-022).

### 2.10 Cell proliferation analysis

Cardiac fibroblasts were cultured in T25 culture flasks (2.0×10^5^ cells/flask, 25 cm^2^ growth area) for each treatment group. Cells were harvested after trypsinization and seeded on gelatin-coated 96-well plates (1×10^4^ cells/well). Fibroblasts were treated with serum-deprived medium for 24 h under Hcy stimulation at 0, 50, 200, 500, 1000 μmol/L concentrations of Hcy. Next, each well was incubated with cell counting kit-8 (CCK-8) solution (100μl) for 2 h. The absorbance was read using a spectrophotometer.

### 2.11 Protein-protein interaction

The gene encoding TRPC3-N (1-369aa) and TRPC3-C (659-836aa) was synthesized by Detai Biologics Co., Ltd. (Nanjing, China). HEK 293-T cells were transfected with the above expression constructs to produce and purify recombinant protein for validating an interaction between TRPC3 and SIRT1 by bio-layer interferometry (BLI).

Cells or cells transfected with indicated vectors were solubilized in cell lysis buffer for IP (Beyotime, China)^[18]^ with proteases and phosphatase inhibitors (pH 7.4). Pre-cleared cell lysates were incubated with equal amounts of primary antibodies (2-5 μl) or IgG at 4°C before performing the pull-down with 50 μl of 1:1 Protein A/G PLUS-Agarose (Santa Cruz Biotechnology, sc-2003) for 2 h. Beads were washed 4 times with lysis buffer, boiled and elution collected for WB analysis ^[19]^.

### 2.12 Kinetic binding analysis by BLI

BLI experiments were performed using an Octet K2 system (Pall Fortebio Corp, Menlo Park, CA). Recombinant SIRT1 was immobilized on the Anti-Penta-HIS (HIS1K) Biosensors (ForteBio, part no. 18-5120) using black polypropylene microplates (Greiner Bio-One part no.655209). Different concentrations of the TRPC3-C or -N peptide were applied in the mobile phase and the association between the immobilized and flowing proteins was detected. The reference buffer was PBST and 5% DMSO (pH 7.4 PBS, tween 20 0.05% and 5% DMSO v/v). Statistical analysis was performed using Data Analysis 9.0 software. The dissociation rate constant (K_D_) was obtained by curve fitting of the association and dissociation phases of the sensograms.

### 2.13 Protein isolation and western blot

Protein was extracted from heart tissues using a Tissue Homogenizer (PD500-TP12) (Prima, GBR) with RIPA Lysis and Extraction Buffer and protease inhibitor (Thermo Fisher Scientific, MA). Protein concentrations were measured using a BCA protein assay kit (Bio-Rad, CA), and 6-8% SDS-polyacrylamide gel electrophoresis (PAGE) was applied to separate high-molecular-weight proteins (Solarbio, China). Collagen-I antibodies were purchased from Abcam (Cambridge, UK; Col-I-ab6308). TRPC3, SIRT1 and TGF-ß antibodies were purchased from CST (Cell Signaling Technology, Boston, USA; TRPC3-77934, SIRT1-8469, and TGF-ß-3711). The GAPDH antibody was obtained from Proteintech (Chicago, USA; GAPDH-60004-1-Ig). Primary antibodies (1:500) were incubated overnight with 5% bovine serum albumin at 4°C and secondary antibodies (HRP-conjugated, 1:5,000) were incubated for 60 min with TBST at room temperature. Proteins were detected with WESTAR ETA C ULTRA 2.0 (Cyanagen, ITA) and images were evaluated using a ChemiDoc MP imaging system (Bio-Rad, CA).

### 2.14 Quantitative reverse transcriptase polymerase chain reaction (RT-qPCR)

Heart tissues were stored in Allprotect Tissue Reagent (Qiagen, Hilden, Germany) for RT-qPCR. Total RNA was isolated from heart tissue ground over liquid nitrogen and extracted using the standard TRIzol method (Invitrogen, MA). cDNA was synthesized with 2 μg of RNA using the PrimeScript™ 1st Strand cDNA Synthesis Kit (Takara Biomedical Technology, Dalian, China). Primers for TRPC3, SIRT1, TGF-ß, Col-I, GAPDH were obtained from Sangon Biotech (Shanghai, China). RT-qPCR was performed using SYBR^®^ Premix Ex Taq™ (Tli RNaseH Plus) (Takara Biomedical Technology, Dalian, China) under the following conditions, step one: 95°C for 5 min, 95°C for 10 sec (reverse transcription); step two: 40 cycles of 95°C for 5 sec and 60°C for 15 sec (PCR); step three: dissociation protocol. Each cycle threshold (CT) value was normalized to the CT value of GAPDH. Fold changes were calculated with 2^(-ddCT) compared to the control group. Specific primers were designed to amplify SIRT1 (SIRT1-F: 5’ TGTGGTGAAGATCTATGGAGGC 3’ and SIRT1-R: 5’ TGTACTTGCTGCAGACGTGGTA 3’), TRPC3 (TRPC3-F: 5’ CAGTGATGCGAGAGAAGGGT 3’ and TRPC3-R: 5’ CGTAGGCGTAGAAGTCGTCG 3’) and TGF-ß (TGF-ß-F: 5’-CCACCTGCAAGACCATCGAC 3’ and TGF-ß-R: 5’-CTGGCGAGCCTTAGTTTGGAC 3’). All data were normalized to the mRNA levels of the housekeeping gene GAPDH (GAPDH-F: 5’ GGTTGTCTCCTGCGACTTCA 3’ and GAPDH-R: 5’ TGGTCCAGGGTTTCTTACTCC 3’)

### 2.15 Statistical analysis

Continuous variables are shown as the mean±standard deviation. Data were evaluated for normal distribution by the Kolmogorov-Smirnov test, and homogeneity of variance was assessed by Levene’s test. An unpaired Student’s t-test was applied to compare normally distributed variables. Variables with non-normal distribution were compared with the Wilcoxon rank sum test. Fisher’s exact test was used to estimate categorical variables. When the variance was not homogeneous, the Tukey test was applied to test the means of multiple samples. Multiple t-tests were performed for multiple-group comparisons. Student t-tests were used to compare only 2 groups. The same group of objects was exposed to multiple interventions, which was necessarily measured by repeated analysis. SPSS 22.0 software was used for the analyses.

## 3. Results

### 3.1 Patients with AF are more often diagnosed with severe cardiac fibrosis in the atrium than SR patients

As shown in Fig. 1A-C, our clinical experiment demonstrated that the left atrial appendage of AF patients exhibited an increase in TRPC3 and pro-fibrosis proteins, such as TGF-ß and collagen-I (Col-I). This was accompanied by a down-regulation of SIRT1 was detected at both the protein and mRNA levels. Fig. 1D-F shows that patients with AF more often presented with atrial fibrosis compared to patients with SR, as detected by IHC and Masson staining experiments. In addition, we observed by electron microscopy that the ultra-structure of atrial tissues was severely damaged in AF patients compared with SR patients (Fig. 1G). For example, the fascicle was ruptured, the crude and fine filaments were loosely arranged, and the mitochondria displayed compensatory enlargement, hyperplasia, and disordered crista. The baseline characteristics of AF and SR patients are presented in Table 1, which shows that the morbidity of hyperhomocysteinemia and the QTc prolongation in AF patients exceeded that in SR patients. On the other hand, the incidence of hypertension was similar in AF and SR patients. Intriguingly, the phenomenon of AF combined with HF was commonly observed, accounting for 33.3% of patients with AF, which is consistent with the current statistics of this comorbidity.

**Figure 1.**
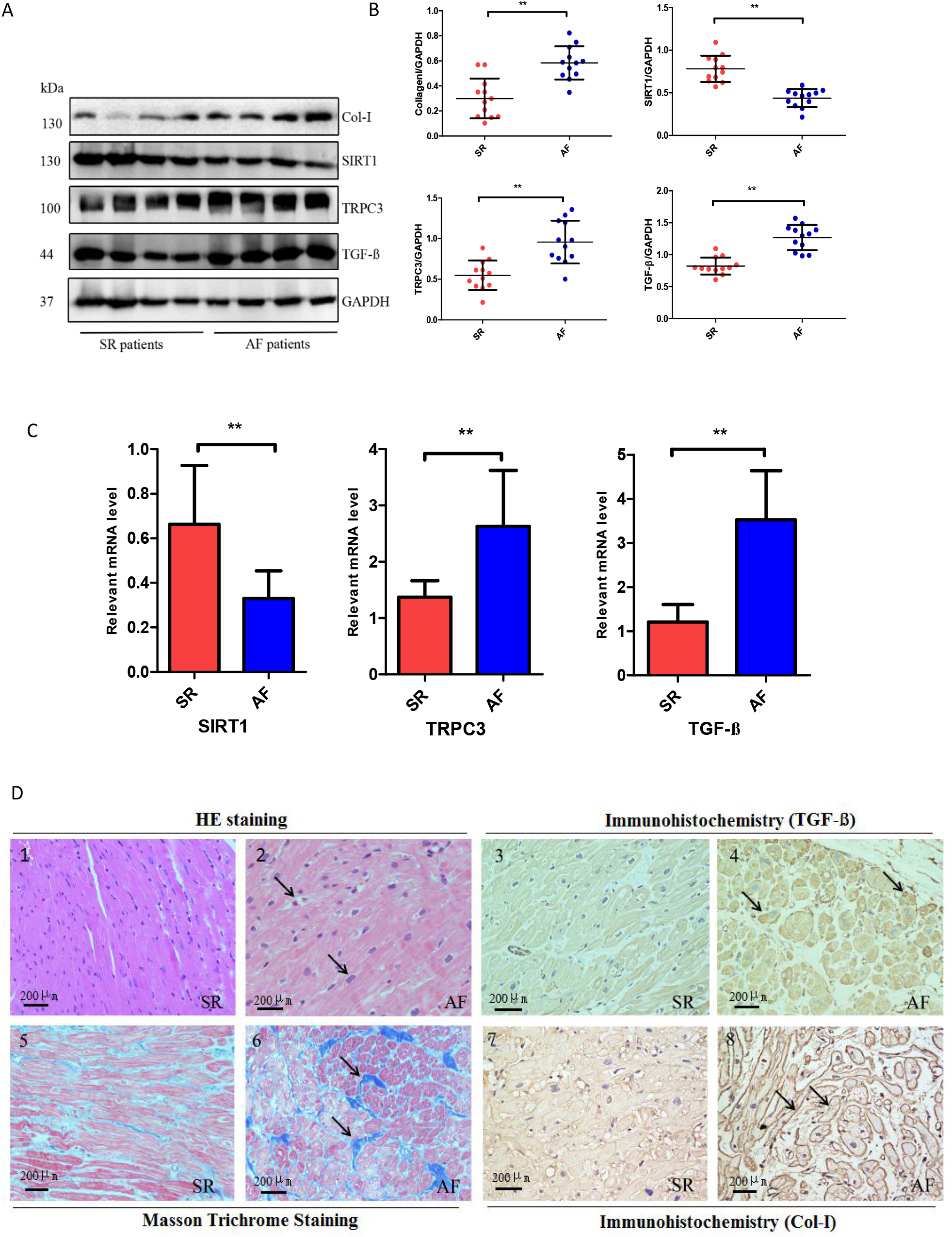
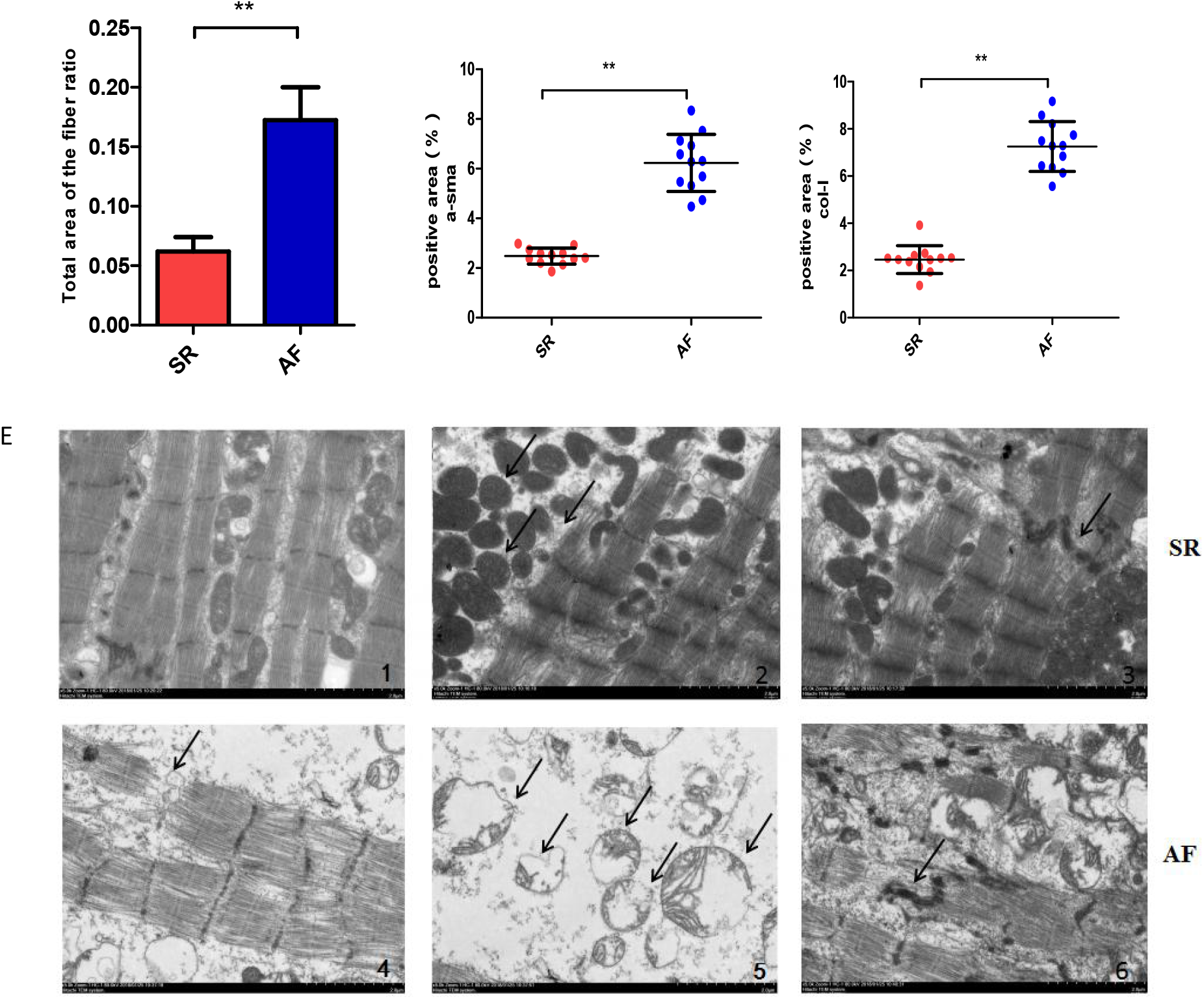
High plasma levels of Hcy are closely related to the occurrence of cardiac fibrosis. **A-B.** The expression levels of TRPC3 and the relevant proteins, such as TGF-ß and Collagen-I in AF patients. C. The relative mRNA level. **D1-D2.** The atrial nucleus in patients with AF is showen with hematoxylin and eosin (HE) staining, Masson’s trichrome-stain and immunohistochemistry, as well as statistical results of positive areas and ratio to total area of the fibre (Magnification × 100, Scale bar=200 μm; n=4 per group). **E.** The ultra-structure of atrial tissues was observed by electron microscopy: **E1-E3:** SR patients, **E4-E6:** AF patients. Error line indicates mean and standard deviation. * denotes p<0.01, ** denotes p<0.001. Multiple t-tests were performed using step-down bootstrap sampling to control the family-wise error rate at 0.05 for 1B, C, and unpaired t-test was used for 1D. Fisher’-s exact test was used for analysing Table 1.

**Table 1.**
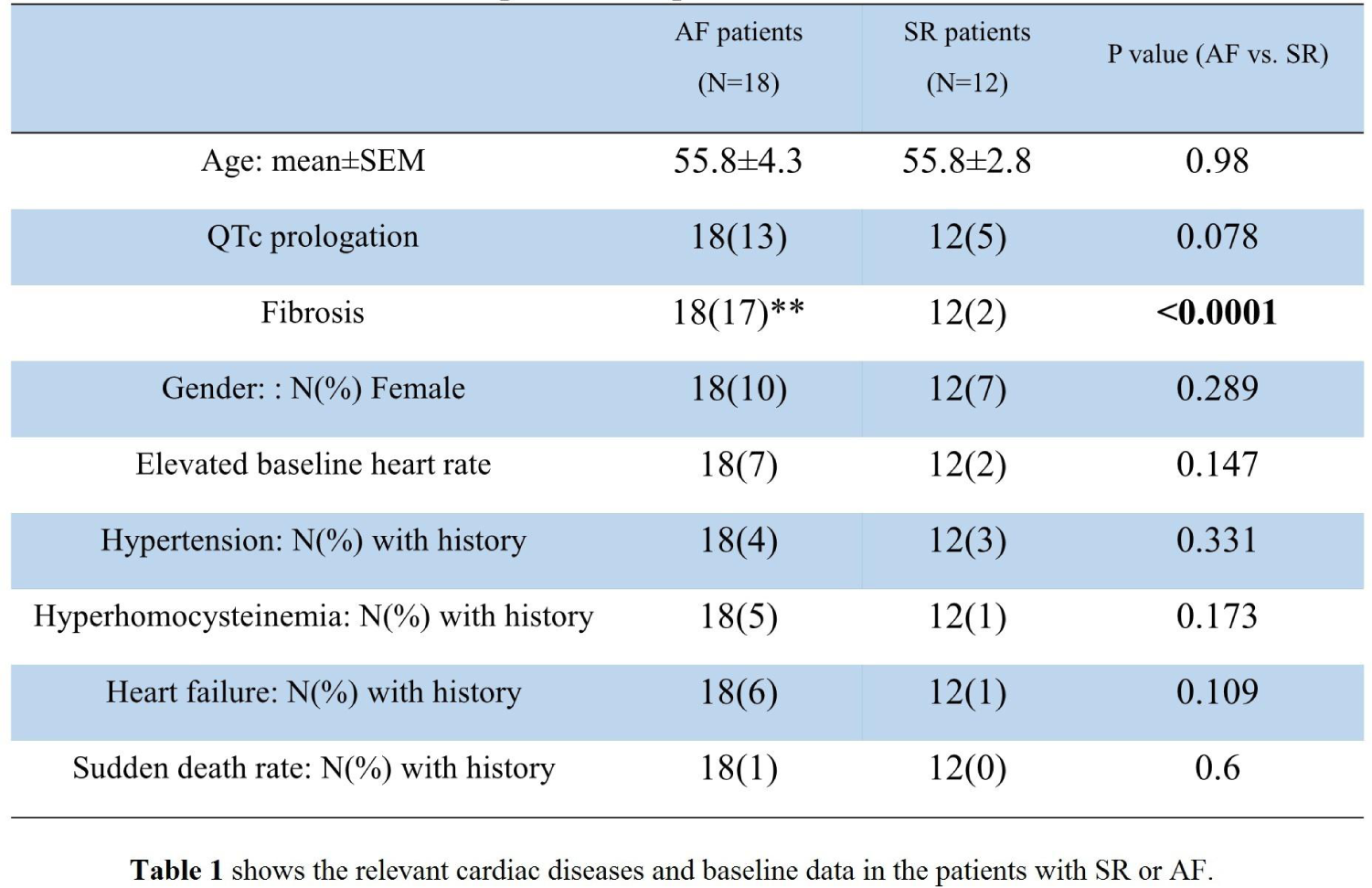
Comparison the patients between AF and SR **Table 1** shows the relevant cardiac diseases and baseline data in the patients with SR or AF.

|  | AF patients<br>(N=18) | SR patients<br>(N=12) | P value (AF vs. SR) |
| --- | --- | --- | --- |
| Age: mean±SEM | 55.8±4.3 | 55.8±2.8 | 0.98 |
| QTc prologation | 18(13) | 12(5) | 0.078 |
| Fibrosis | 18(17)** | 12(2) | **<0.0001** |
| Gender: : N(%) Female | 18(10) | 12(7) | 0.289 |
| Elevated baseline heart rate | 18(7) | 12(2) | 0.147 |
| Hypertension: N(%) with history | 18(4) | 12(3) | 0.331 |
| Hyperhomocysteinemia: N(%) with history | 18(5) | 12(1) | 0.173 |
| Heart failure: N(%) with history | 18(6) | 12(1) | 0.109 |
| Sudden death rate: N(%) with history | 18(1) | 12(0) | 0.6 |

### 3.2 Development of HFpEF in high-Hcy fed mice with TAC

Fig. 2A shows how the Hcy diet was administered to transverse aortic constriction (TAC) mice (n=25) at 4-16 weeks. TAC mice that received the high-Hcy diet were classed as the HH group and those that received a normal diet as the NH group, with mice undergoing sham surgery regarded as the SH group. To induce AF, we administered a constant dose of acetylcholine (0.5 μl/mg), which was injected into each mouse through the caudal vein. Fig. 2B-E reveals echocardiographic results at 4, 7 and 16 weeks of age. The ejection fraction (EF) was not significantly different at baseline (4 weeks) among the three groups (Fig. 2B). The interventricular septal thickness (IVS-S) in HH mice was markedly greater than that in the NH mice at 16 weeks of age (1.914±0.167 in HH vs 1.832±0.134 in NH, p<0.001) (Fig. 2B). The results confirmed that baseline E/A ratios were not different among SH, NH and HH mice at 4 weeks of age before TAC surgery, whereas the E/A ratio was decreased in HH mice compared with the NH mice at 16 weeks, although no statistically significant results were observed at 7 weeks (1.288±0.085 in NH vs. 1.255±0.128 in HH, p>0.05; Fig. 2B), with greater significant differences in the values among the NH and HH groups (1.099±0.101 in HH and vs. 1.205±0.083 in NH, p<0.001; Fig. 2B), indicating impaired diastolic function. The ratio of transmitral flow velocity to annular velocity (E/E’) is a non-invasive parameter for left ventricular (LV) stiffness ^[20]^. There was a slight variability in the E/E’ ratio at the baseline at 4 weeks, and the value of E/E’ was increased in HH mice beyond that in the NH mice at 16 weeks (21.89±3.738 in HH vs. 19.53±3.206 in NH, p<0.001 at 16 weeks but 16.82±2.495 in HH vs. 15.97±2.513 in NH, p>0.05 at 7 weeks; Fig. 2B). The measurements of LVIDd were markedly increased following TAC (NH and HH) was observed, especially in the HH group (Fig. 2B, Table2). Evolving mitral flow velocity patterns were analysed during different time periods (Fig. 2C). The ultrasonic images at different periods were obtained from a longitudinal section of the heart (Fig. 2D). In addition, increasing posterior wall thickness at end-diastole (PWTd) and posterior wall thickness at end-systole (PWTs) measurements were concurrently noted (Table 2). However, the heart rate (reported here as the RR interval) was not different among the three groups (Table 2).

**Figure 2.**
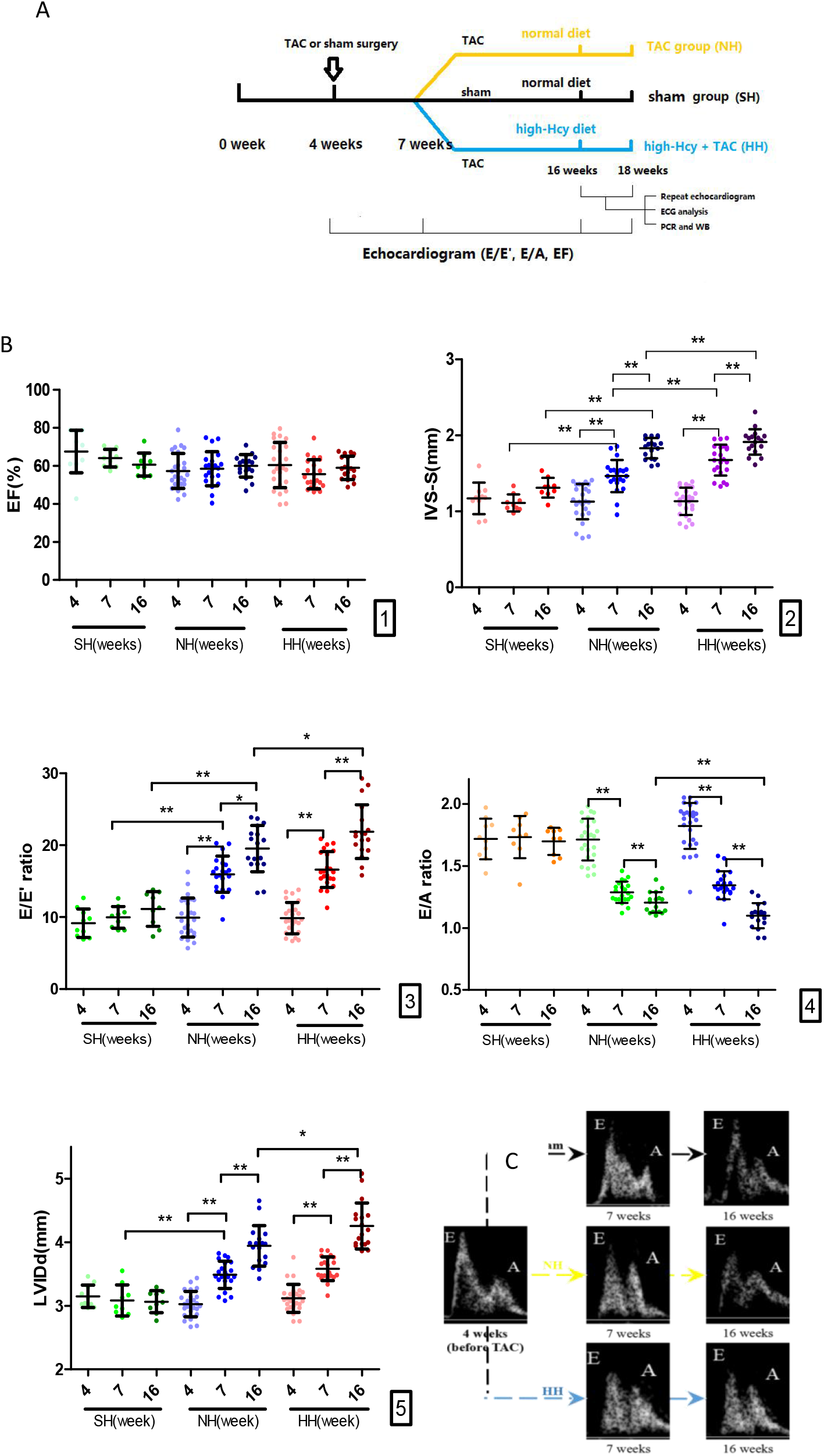
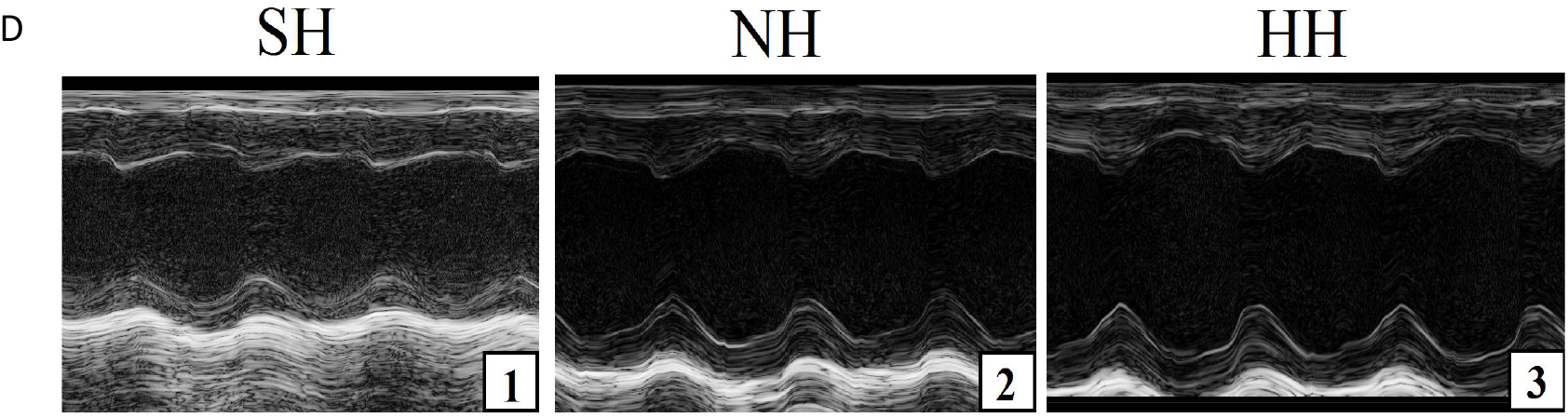
Timeline for establishing mouse HFpHR model with a high Hcy diet and echocardiographic parameters for assessing systolic and diastolic function. **A.** Mice underwent TAC or sham surgery at 4 weeks of age in the HH group (n=25), which were fed a high-Hcy diet (4% methionine) from 4-18 weeks of age. The NH group (n=25) also underwent the above TAC operation but were fed a normal diet. Mice that underwent sham surgery and were fed a normal diet were considered the sham (SH) group (n=10). Repeat echocardiogram (ECG) was performed at 16 and 18 weeks of age. We performed programmed ECG analysis, RT-qPCR and western blot in the three groups at 18 weeks of age. **B1.** The value of ejection fraction (EF) in SH, NH and HH mice. **B2.** The value of inter-ventricular septum thickness (IVS-S). **B3, B4.** Baseline E (early filling) /A (atrial filling) and E/E’ ratios among the three groups. **B5.** The value of left ventricular internal diastolic diameter (LVIDd). **C.** Representative images from pulsed-wave Doppler showing E and A wave. **D.** Representative M-mode images of the three groups. Error line indicates mean and standard deviation. * denotes p<0.01 and ** denotes p<0.001. Mixed model regression with post-hoc testing (Tukey adjustment) was used for 2B.

**Table 2.** Echocardiography Parameters and Heart Mass Ratio

|  | SH |  |  | NH |  |  | HH |  |  |
| --- | --- | --- | --- | --- | --- | --- | --- | --- | --- |
| Weeks | 4 | 7 | 16 | 4 | 7 | 16 | 4 | 7 | 16 |
| N | 10 | 9 | 9 | 25 | 21 | 18 | 25 | 22 | 17 |
| Heart rate | 489±32 | 475±26 | 447±38 | 458±30 | 451±29 | 439±27 | 453±36 | 479±33 | 451±37 |
| E wave | 331.8±36.95 | 345.5±19.31 | 339.2±27.95 | 358.9±13.59 | 320.4±17.41 | 300.9±30.21 | 355.4±10.54 | 313.4±18.41 | 280.2±34.72 |
| E’ wave | 37.44±7.48 | 35.48±6.48 | 30.84±6.077 | 38.74±10.77 | 20.57±3.54 | 15.94±3.838 | 37.70±8.015 | 19.23±2.87 | 13.17±2.660 |
| A wave | 194.9±29.31 | 201.2±23.26 | 200.6±21.668 | 211.5±20.54 | 249.9±21.12 | 250.6±29.26 | 197.0±20.65 | 234.5±21.13 | 257.3±42.35 |
| EF% | 67.5±11.2 | 64.0±4.6 | 60.7±6.1 | 57.3±9.2 | 57.7±9.7 | 60.0±5.9 | 60.4±11.9 | 55.6±7.7 | 59.0±6.1 |
| FS% | 41.5±3.5 | 39.7±3.0 | 42.8±2.1 | 39.7±1.6 | 45.3±2.4 | 38.7±4.2 | 39.6±3.8 | 43.2±4.7 | 37.4±2.9 |
| LVIDd, mm | 3.148±0.176 | 3.082±0.245 | 3.064±0.174 | 3.026±0.200 | 3.488±0.214 | 3.943±0.320 | 3.188±0.220 | 3.583±0.187 | 4.256±0.362 |
| LVIDs, mm | 2.03±0.19 | 1.78±0.34 | 2.12±0.09 | 1.58±0.28 | 2.62±0.26 | 3.31±0.39 | 1.76±0.41 | 2.47±0.15 | 3.23±0.21 |
| PWTd, mm | 0.59±0.09 | 0.89±0.15 | 0.99±0.12 | 0.67±0.10 | 0.79±0.18 | 0.93±0.11 | 0.71±0.18 | 0.85±0.06 | 1.21±0.21 |
| PWTs,mm | 0.78±0.12 | 0.89±0.17 | 1.03±0.09 | 0.83±0.23 | 0.94±0.19 | 1.27±0.14 | 0.87±0.11 | 1.09±0.16 | 1.43±0.34 |
| BNP,pg/ml |  |  | 48±6 |  |  | 224±28 |  |  | 293±47 |
| Heart weight, g |  |  | 0.45±0.054 |  |  | 0.56±0.11 |  |  | 0.63±0.10 |

**Table 3.** Animal groups.

| Group | Surgery | Diet | Intraperitoneal injection<br>(20mg/kg/day) |
| --- | --- | --- | --- |
| SH | Sham | Normal |  |
| NH | TAC | Normal |  |
| HH | TAC | High-Hcy |  |
| HH+saline | TAC | High-Hcy | Saline |
| HH+Res | TAC | High-Hcy | Resveratrol (Res) |
| HH+FC | TAC | Folic acid+High Hcy | Saline |

### 3.3 Arrhythmia induced by acetylcholine and atrial fibrosis in mice

Fig. 3A-C shows the results of ECGs recorded in the three groups within five minutes of acetylcholine injection. Normal SR was typically observed in the ECGs of the SH group (n=7/9), and the mean speed was 125 ms (Fig. 3A). ECGs indicating AF include the following three characteristics: irregular small F waves, no P wave, and an unequal RR interval, and were frequently observed in the HH group (n=10/17) (Fig. 3B). In addition, the features of ventricular fibrillation (VF) include the disappearance of the QRS wave, which is replaced by different ventricular fibrillation waves, and was observed in the NH (n=3/18) and HH (n=4/17) groups (Fig. 3C, 3D). Fig. 3D illustrates that AF was induced in 10 out of 17 (58.8%) mice in the HH group; the frequency of AF was higher in the HH group than in the NH group [8 of 18 (44.4%)], but the difference was not statistically significant. However, both the HH and NH groups exhibited significantly higher induction ratios of AF than the SH group that did not undergo TAC (2 of 9 (22.2%)) (P<0.01). VF was not observed in the SH mice, but monomorphic VF was observed in the NH [3 of 18 (16.7%)] and HH [4 of 17 (23.5%)] groups (Fig. 3D). Despite the lack of statistical significance, the propensity to develop AF with or without feeding a high-Hcy diet was higher in groups that underwent TAC (HH and NH groups) than in the SH group (Fig. 3D). Interestingly, the duration of induced AF was longer in both the HH and NH groups than in the SH group, especially in the HH group (Fig. 3E).

**Figure 3.**
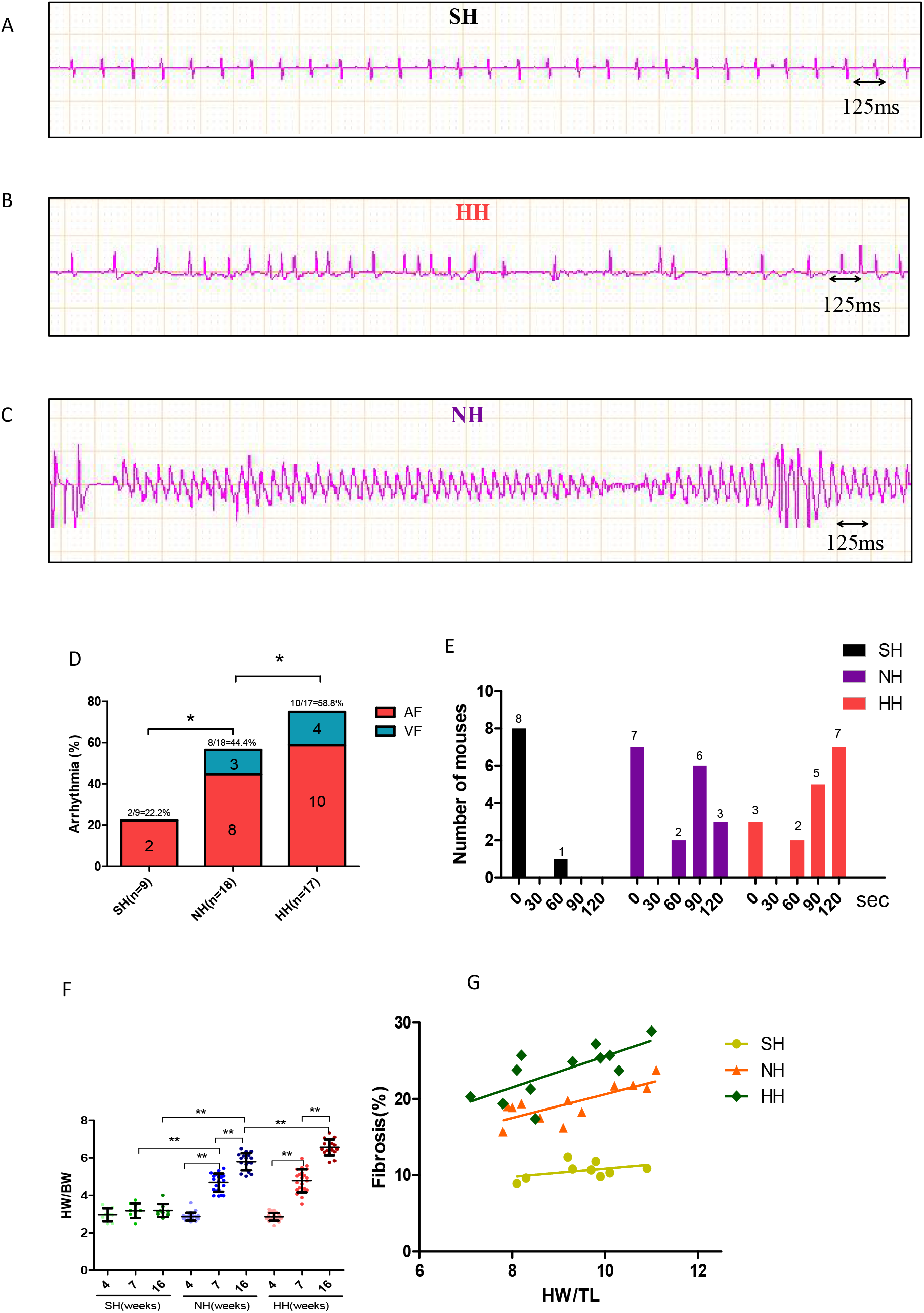
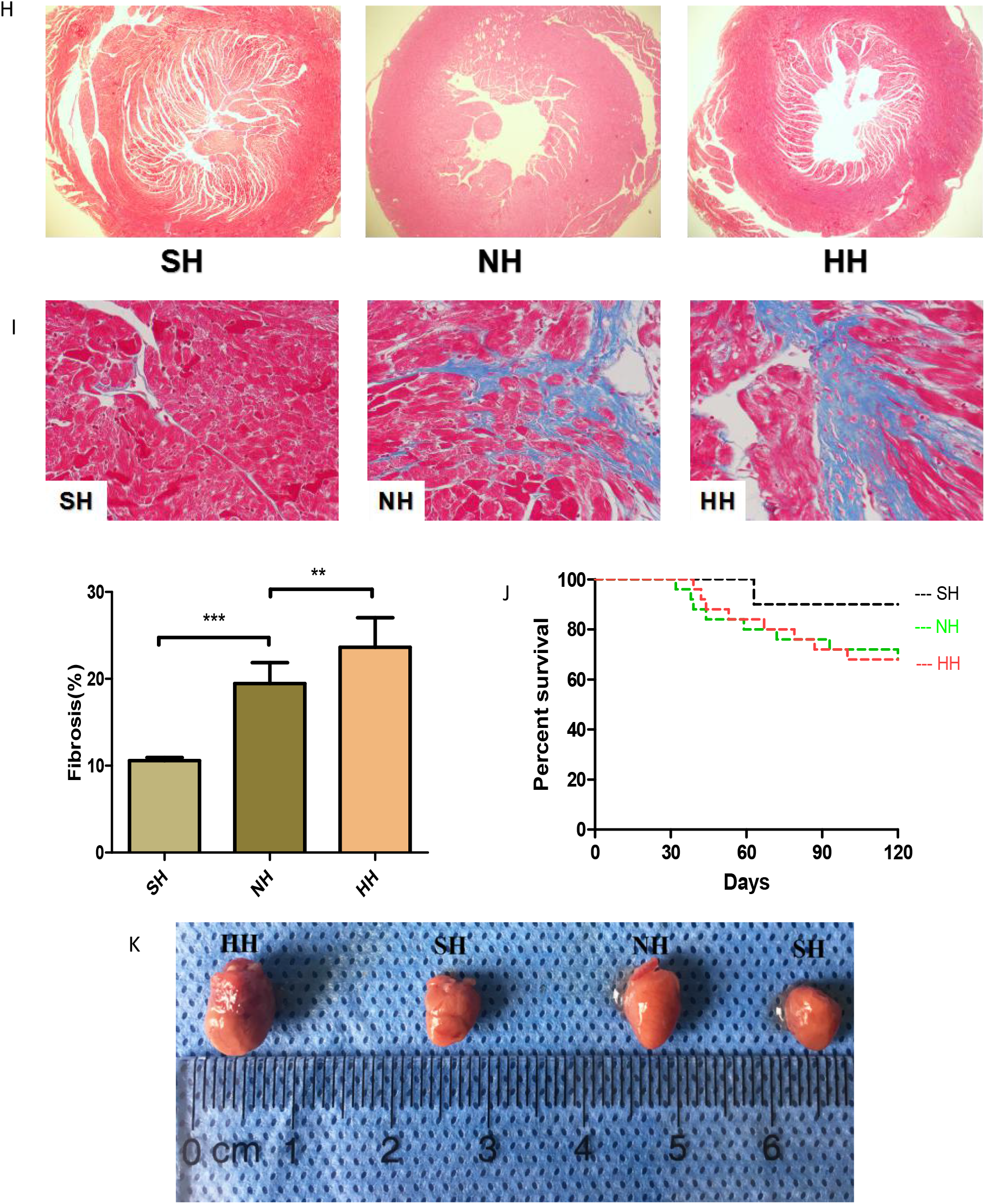
Programmed acetylcholine stimulation (PES) and ECG analyses in SH, NH and HH mice. **A.** Normal SR was typically observed in the ECGs of the SH group (n=7/9), and the speed was 125 ms. **B.** ECGs indicating AF were frequently observed in the HH group (n=10/17). **C.** Ventricular fibrillation (VF) was observed in the NH (n=3/18) and HH (n=4/17) groups. **D.** Increased susceptibility to AF in the HH group (N=10/17) than in the NH group (N=8/18), with the lowest susceptibility in the SH group (N=2/9). **E.** The duration of AF in the three groups. **F.** The value of heart weight-to-body weight (HW/BW). **G.** The heart weight-to-tibia length ratio (HW/TL). **H.** Representative Masson staining showed transverse section in atria. **I.** Representative Masson staining images with magnified local images in atrial tissues. **J.** Survival rate of the three groups over different time periods. **K.** Results are represented as cardiac sizes in the three groups. * denotes p<0.05, ** denotes P<0.01. Mixed model regression with post-hoc testing (Tukey adjustment) was used in 3F, 3G, and Fisher’s exact test was used in 3D.

Next, we studied whether Hcy affects the process of atrial fibrosis in animal experiments. There was a slight fluctuation in the heart weight/body weight ratio (HW/BW) at the baseline at 4 weeks, and the value of HW/BW was increased in HH mice beyond that in the NH mice at at 16 weeks (6.552±0.418 in HH vs 5.795±0.461 in NH, p<0.001 at 16 weeks but 4.777±0.610 in HH vs 4.679±0.479 in NH, p>0.05 at 7 weeks; Fig. 3F), similar changes were observed for the heart weight-to-tibia length ratio (HW/TL) at 16 weeks of age (Fig. 3G). In addition, Masson’s trichrome staining simultaneously affirmed that the HH groups were more vulnerable to fibrosis, compared to the NH group (Fig. 3I). Moreover, Transverse sections of atrium showed a greater enlargement in the HH group, compared with the NH group (Fig. 3H). and further demonstrated that Hcy could ameliorate the adverse influence of TAC-induced HF. Survival analysis showed that mice in the NH and HH groups exhibited higher mortality, especially after 10 weeks, than that in the SH group, as shown in Fig. 3J.

### 3.4 SIRT1-overexpression and TRPC3-KD mice can efficiently control Hcy-mediated atrial fibrosis

Fig. 4A shows the procedure for intravenously injecting purified SIRT1-overexpression or TRPC3-shRNA lentivirus through the tail vein of mice at one week of age, respectively regarded as SIRT1-overexpression or TRPC3-KD mice. Scramble lentivirus was injected into vector mice as a negative control. At 4 weeks, we performed the TAC operation and fed mice with a high-Hcy diet for 10 weeks. Next, echocardiography was used to detect the EF and other indicators at 7 and 16 weeks. Fig. 4B-C shows that the degree of fibrosis was increased in the HH+vector group compared with the NH+vector group (Fig. 4B1). Interestingly, when TAC mice were fed a normal diet, the HW/TL ratio was lower in the TRPC3-KD or SIRT1-overexpression mice, which indicates significant suppression of the extent of atrial fibrosis compared to the NH+vector mice (Fig. 4B3). Although the degree of fibrosis was also decreased in the TRPC3-KD or SIRT1-overexpression mice with a high-Hcy diet, than in the HH+vector mice (Fig. 4B2). There was also a slight increase in the level of atrial fibrosis in the TRPC3-KD or SIRT1-overexpression mice with a high-Hcy diet, compared to those mice with a normal diet (Fig. 4B4). According to their echocardiography, the IVS-S at end-systole and LVIDd were efficiently alleviated by the above treatments (Supplement Fig. 1A). Meanwhile, the Masson staining of TRPC3-KD+HH and SIRT1-overexpression+HH mice was dramatically decreased in comparison with the HH mice (Fig. 4C), whilst the EF level remained at a relatively stable state (Supplement Fig. 1B). In addition, transgenic mice displayed no apparent changes in cardiac structure and function (Supplement 1C, 1D). This result confirmed that despite the influence of the high-Hcy diet and the TAC surgery, the protein level of TRPC3 and the related TGF-ß signalling pathway ^[9]^ were inhibited in the TRPC3-KD+HH group, while SIRT1 was increased. Similar results were observed in the SIRT1-overexpression+HH group with the higher level of SIRT1 (Fig. 4D). In general, the TRPC3-KD and SIRT1-overexpression mouse models generated by injecting purified lentiviruses exhibited dramatically abrogated Hcy-induced fibrosis in atrial fibroblasts and decreased levels of pro-fibrosis proteins, such as TGF-ß and Col-I (p<0.001). Fig. 4E shows the procedure for intraperitoneal injection of Res 20mg/kg/d for 21 days from the fourth week, whilst another group were fed a diet containing folic acid and were injected with the same dose of saline, respectively regarded as HH+Res or HH+FC mice. Interestingly, regardless of whether folic acid and vitamins were added to the food or not, there was no notable protection from the damage caused by high homocysteine levels (Fig. 4F-4I). However, Res attenuated the enhancement of TRPC3 and decrease in SIRT1 induced by the high-Hcy diet combined with TAC and controlled the TGF-ß improvement (Fig. 4F-4H). In addition, the mRNA levels of TRPC3 and SIRT1 in the above groups were consistent with their protein levels (Fig. 4I).

**Figure 4.**
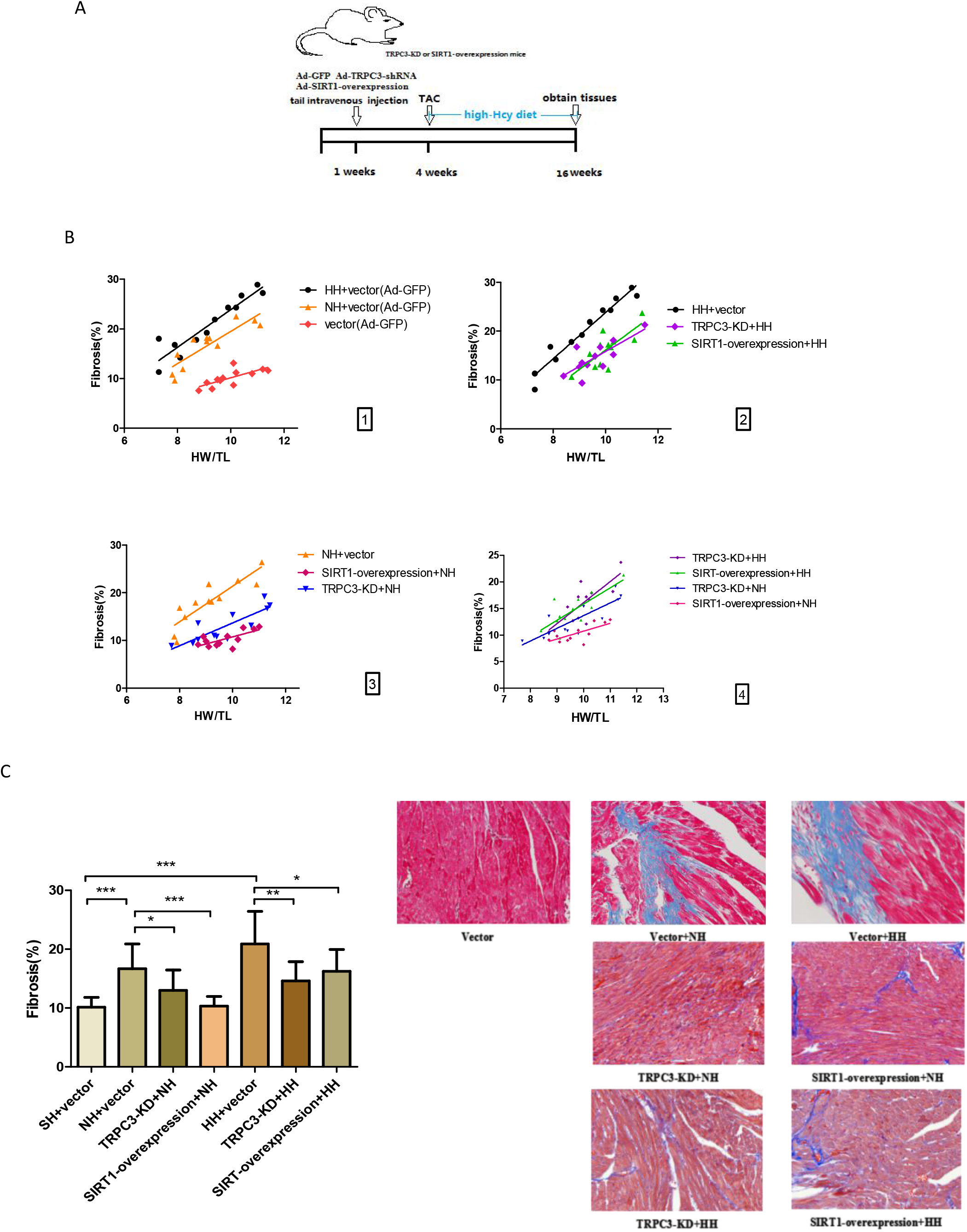
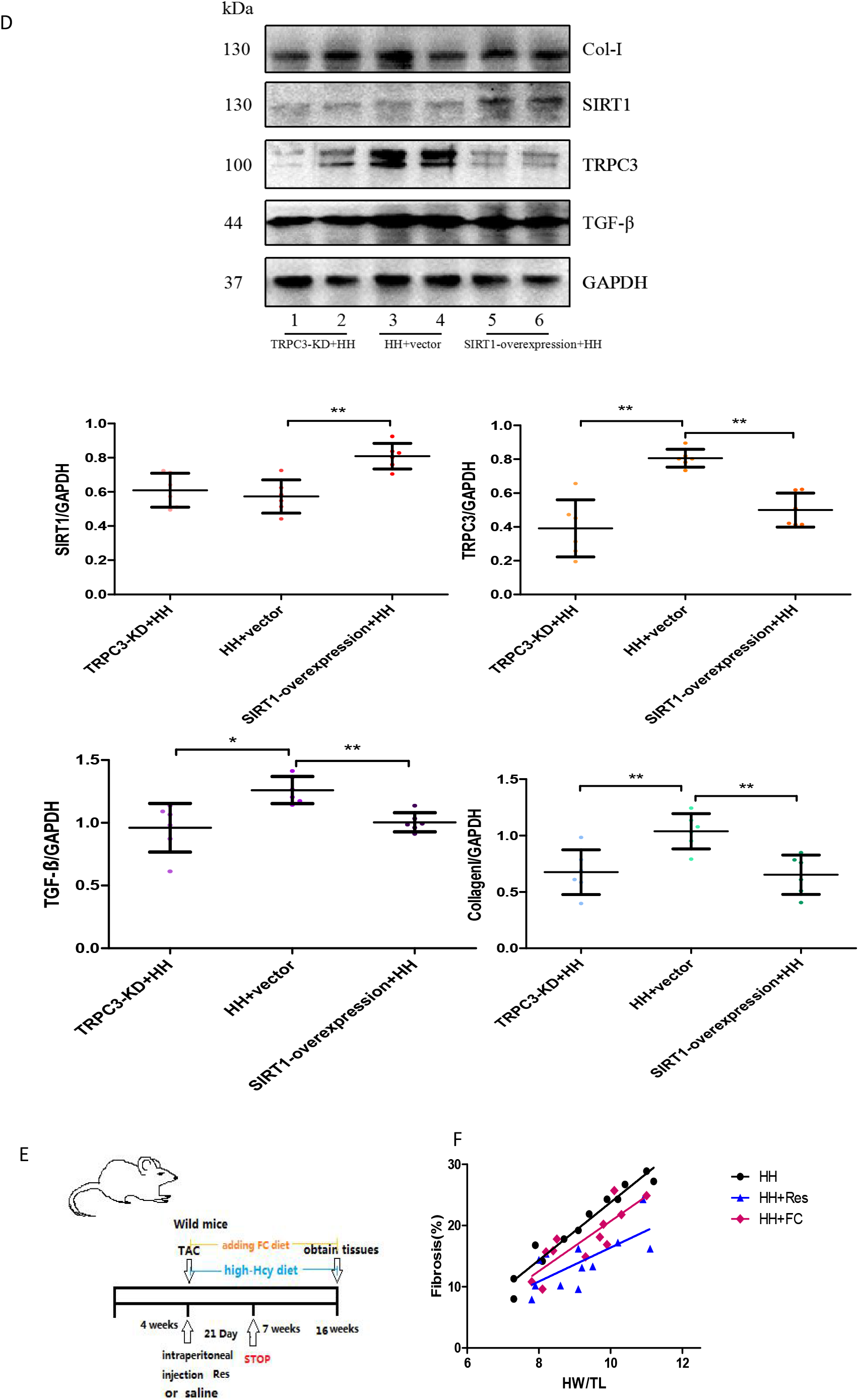
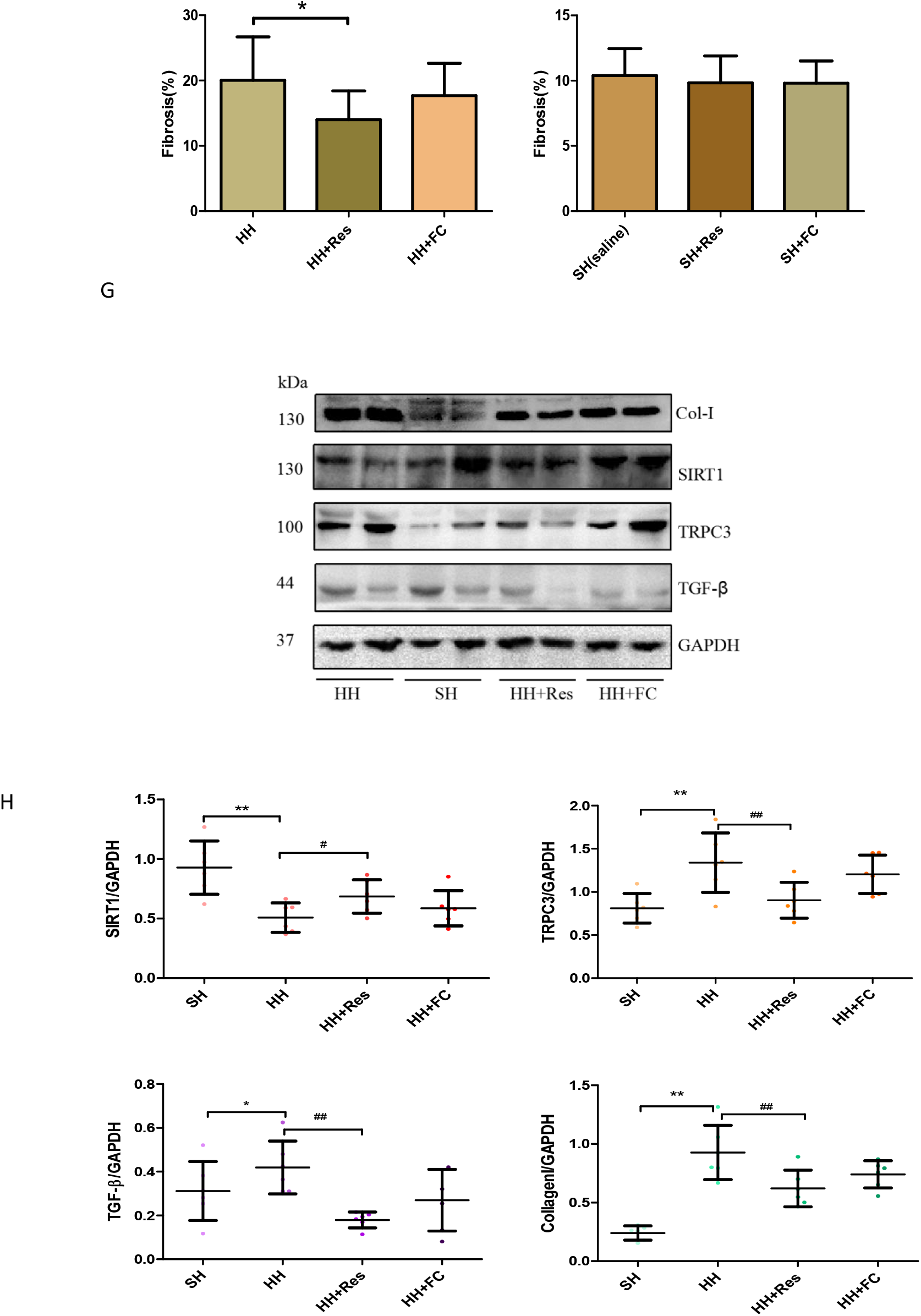
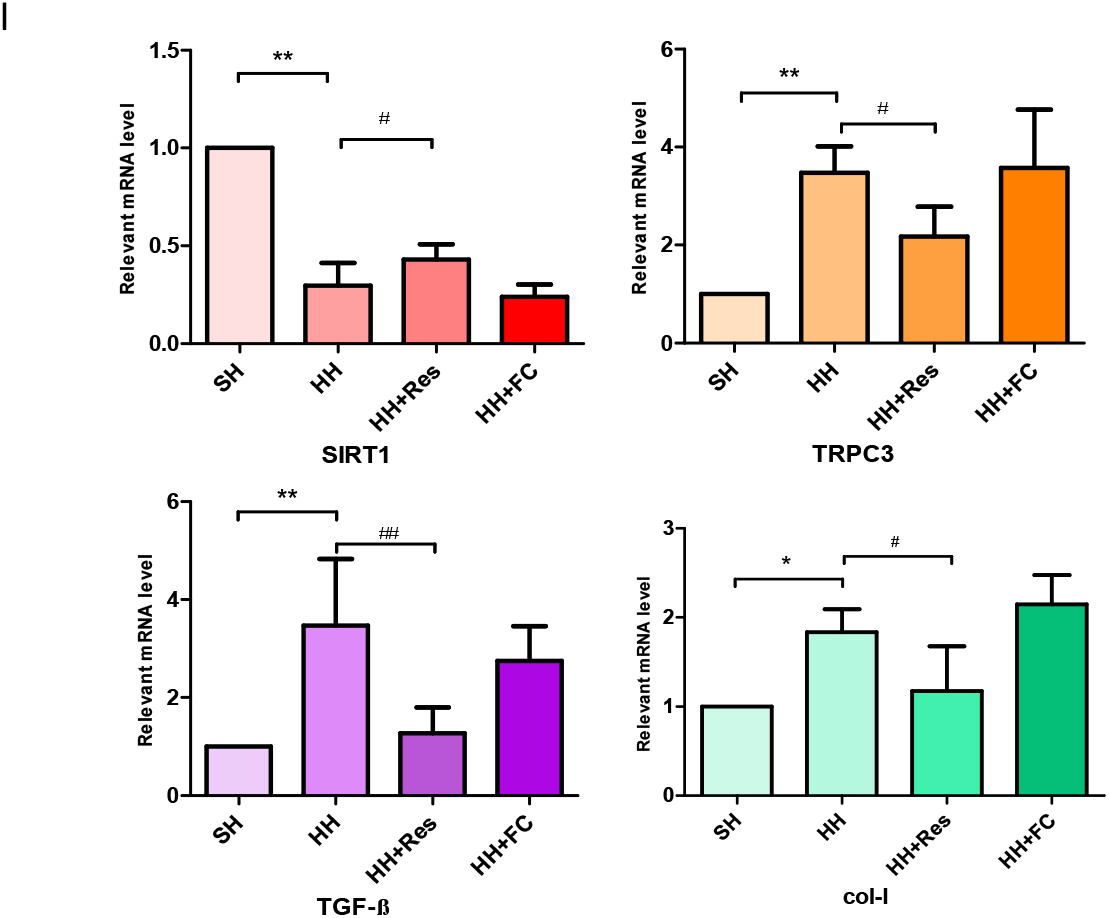
TRPC3-KD or SIRT1 overexpression in mice could effectively alleviate atrial fibrosis. **A.** C57B6 mice (n=10) at 1 week of age were injected with purified lentivirus via the caudal vein. Mice underwent TAC and were fed with a high-Hcy diet at 4 weeks of age, with relevant tissues obtained after 16 weeks. **B, F.** Heart weight-to-tibia length ratio (HW/TL). **C.** The Masson staining images with magnified local images in atrial tissues. **D.** The protein levels of TRPC3 and TGF-ß measured in the three groups. **E.** Ten C57B6 mice underwent TAC at 4 weeks of age and were fed with a high-Hcy and folic acid mixed diet during 4-16 weeks (HH+FC group). **F.** A subset of these animals were injected with Res (20mg/kg/day) for 21 day (HH+Res group). **G.** The statistical total area of the fibre ratio. **H, I.** The protein and mRNA expression levels of SIRT1 and TRPC3. Error line indicates mean and standard deviation. * denotes P<0.05 and ** denotes P<0.01. Mixed model regression with post-hoc testing (Tukey adjustment) was used for 4B and 4F. Multiple t-tests were performed using step-down bootstrap sampling to control the familywise error rate at 0.05 for 4C, 4D and 4G-4I.

### 3.5 Hcy can promote the proliferation and differentiation of atrial fibroblasts and modulate the relevant protein levels

First, the results revealed that the expression levels of total TRPC3 were significantly increased by the intervention of Hcy in a dose-dependent manner (Fig. 5A) compared with those in the control group (Hcy 0μmol/L). However, those expression levels were significantly decreased by Pyr-10 treatment (p<0.001), which specifically inhibits TRPC3 (Fig. 5B). These results indicated that Hcy could increase cardiac TRPC3 expression in mouse atrial fibroblast, and that this effect was inhibited by Pyr-10. Second, we observed that the protective effects of SIRT1 against cardiac remodelling were decreased by Hcy (p<0.001). Intriguingly, application of the activator of SIRT1 (Res) could prevent the Hcy-induced enhancement of TRPC3 level. In addition, the above effects of Hcy on atrial fibroblast were accompanied by a reduction in the expression level of fibrosis-related proteins, such as TGF-ß, which was demonstrated to be a pivotal molecule in fibrosis (p<0.001) (Fig. 5B). Finally, when the inhibitor of SIRT1 (Salermide) was applied to mice fed a high-Hcy diet, the negative effects on atrial fibroblast were exacerbated (Fig. 5B).

**Figure 5.**
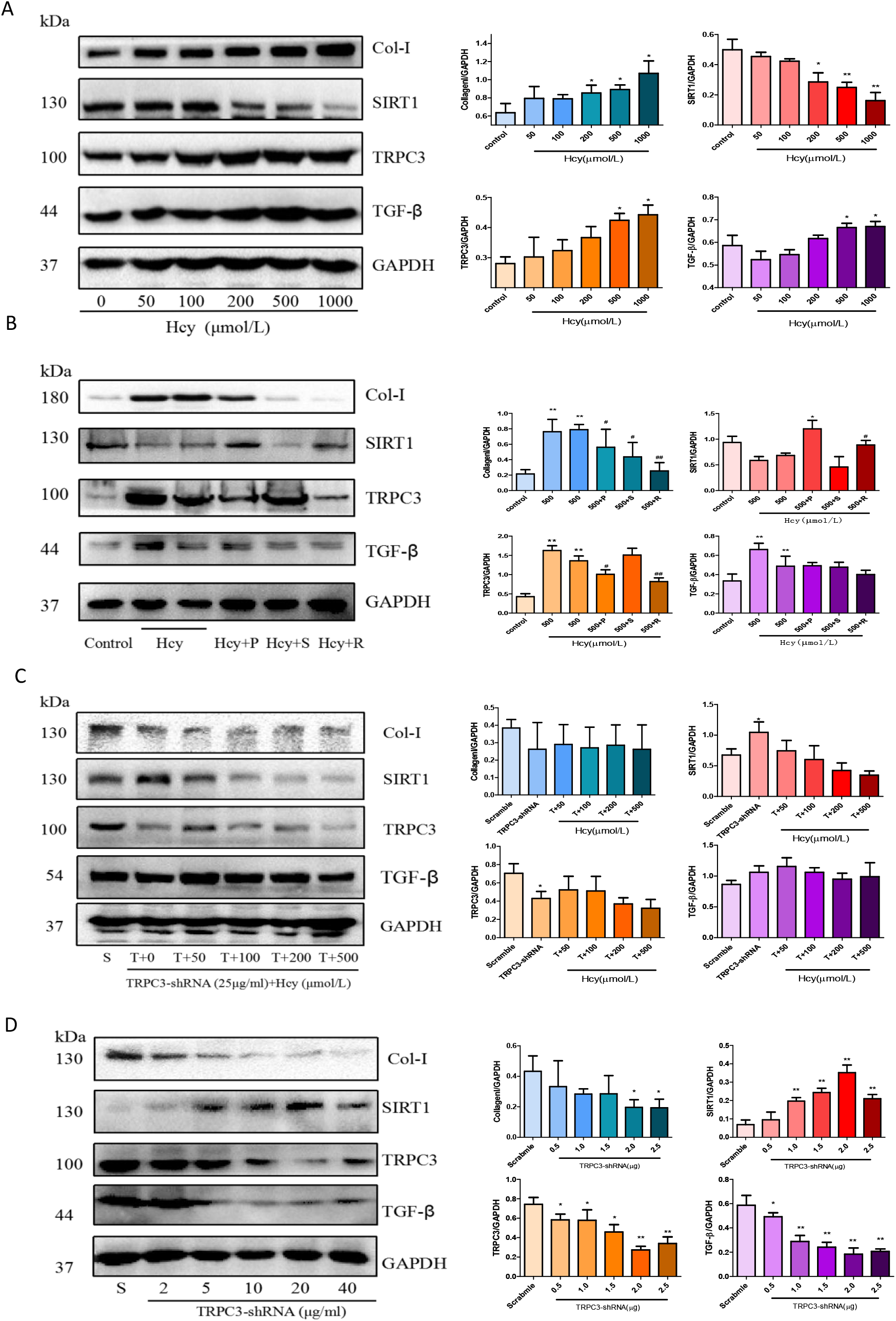
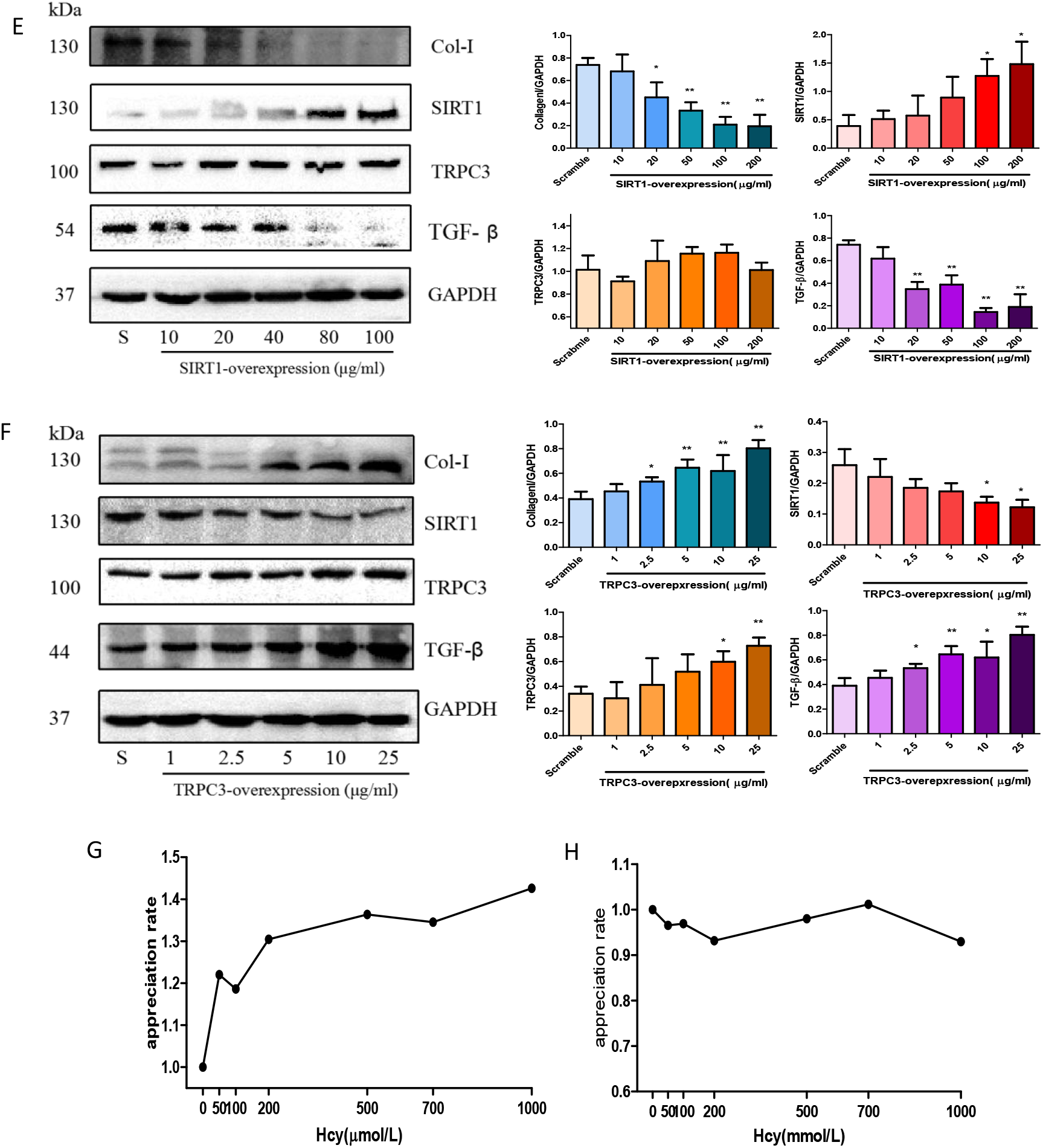
Hcy modulates the expression of TRPC3 protein and affects the activity of SIRT1. **A-B.** The expression levels of TRPC3, SIRT1, TGF-ß and Col-I in mouse atrial fibroblast. **C-D.** The protein levels of TRPC3, SIRT1 and TGF-ß in TRPC3-shRNA-transfected stable cell lines. **E-F.** The expression levels of TRPC3, SIRT1, TGF-ß and Col-I in in TRPC3-overexpression or SIRT1-overexpression stable cell lines. **G, H.** The promotion of fibroblast proliferation was detected by CCK-8. Error line indicates mean and standard deviation. * denotes p<0.05 and ** denotes p<0.01 vs. scramble; # denotes p<0.05, ## denotes p<0.01 vs. (500 μmol/L) Hcy group. Unpaired t-test was used for 5A-5F, and mixed model regression with post-hoc testing (Tukey adjustment) was used for 5G, 5H.

Second, we examined the expression levels of cardiac TRPC3 using purified lentivirus to transfect primary cultured atrial fibroblast. Supplement Fig. 2A shows the fluorescence intensity when the multiplicity of infection (MOI) value was 1, 5, 10 and 20. Of note, the transfection efficiency was highest at 20 MOI. Cell viability was not significantly affected by lentivirus transfection with or without Hcy stimulation (Supplement Fig. 2B). We also detected the protein level of TRPC3 and SIRT1. TRPC3-shRNA-transfected stable cell lines exhibited attenuated Hcy-induced atrial fibrosis, concomitant with lower protein levels of TRPC3 and TGF-ß (Fig. 5C). Moreover, regardless of the transfection of TRPC3-shRNA or SIRT1-overexpression lentivirus, the protein levels of SIRT1 increased as the MOI value increased; the level of TRPC3 decreased in cells transfected by TRPC3-shRNA (Fig. 5D), but not in cells transfected by SIRT1-overexpression lentivirus (Fig. 5E). It is worth emphasizing that the protein level of TGF-ß was decreased even though the level of TRPC3 was not markedly changed in SIRT1-overexpression stable cell lines (Fig. 5E). On the other hand, transfection of TRPC3-overexpression plasmids into HEK293 cells elicited the overexpression of TRPC3 protein as well as the suppression of endogenous SIRT1 protein levels in a dose-dependent manner along with the promotion of TGF-ß expression (Fig. 5F), which suggested that TRPC3 can affect the transcription and translation of SIRT1, which further modulates the protein levels of TGF-ß. Finally, the results suggest that Hcy could promote the proliferation and differentiation of fibroblasts (Fig. 5G). In contrast, the promotion of fibroblast proliferation by Hcy was blocked in TRPC3-shRNA stable cell lines (Fig. 5H).

### 3.6 SIRT1 is an TRPC3-interacting partner

Although TRPC3 is well established to play a central role in transducing cardiac fibrosis signalling ^[21]^, the mechanisms underlying its signal transduction function still remain poorly understood, especially when TRPC3 also regulates TGF-ß. To address this issue, researchers have identified SIRT1 as a modulator of the transcription of TGF-ß-dependent genes, which participate in the process of fibrosis ^[22]^. In a preliminary experiment, we found that Hcy could increase the level of TRPC3; nevertheless, the protein level of SIRT1 was decreased (Fig. 5A). Recent studies have determined that sirtuins can modulate the regulation of a variety of cellular processes associated with RAS. Among them, SIRT1 protects the cell from oxidative stress ^[23]^, indicating that Hcy improves TRPC3 by modulating reactive oxygen species (ROS). SIRT1 could be a potential TRPC3-interacting partner, as both regulate the process of cardiac fibrosis in synergy. To confirm this hypothesis, we performed an in vitro GST pull-down assay using Flag-TRPC3 expressed in HEK293T cells and purified GST-TRPC3 to further validate the TRPC3-SIRT1 interaction in mammalian cells. In addition, co-immunoprecipitation (IP) experiments were performed to corroborate that Flag-TPRC3 and HA-SIRT1 interact in HEK293T cells (Fig. 6B). Endogenous TRPC3 protein immunoprecipitated with a SIRT1 antibody (Fig. 6C), indicating that TPRC3 and SIRT1 could form a protein complex in cells. Moreover, the interaction between endogenous TRPC3 and SIRT1 was markedly induced by Hcy stimulation (Fig. 6D). Finally, immunofluorescence staining showed that TRPC3 and SIRT1 co-localized in the cytoplasm of cells (Fig. 6A), suggesting that the binding of these proteins occurs in the cytoplasm. Next, we determined the functional outcomes of the TRPC3-SIRT1 interaction, and evaluated whether TRPC3 affects the functions of SIRT1. We demonstrated that the regulation of fibrosis by SIRT1 and TRPC3 is antagonistic. We found that the stable cell line transfected with SIRT1-overexpression lentivirus exhibited a low expression level of TRPC3 (Fig. 5K, 5L), which suggested that TRPC3 is required for the activation and translocation of TGF-ß into the nucleus of fibroblasts by mediating the trafficking and activity of SIRT1.

**Figure 6.**
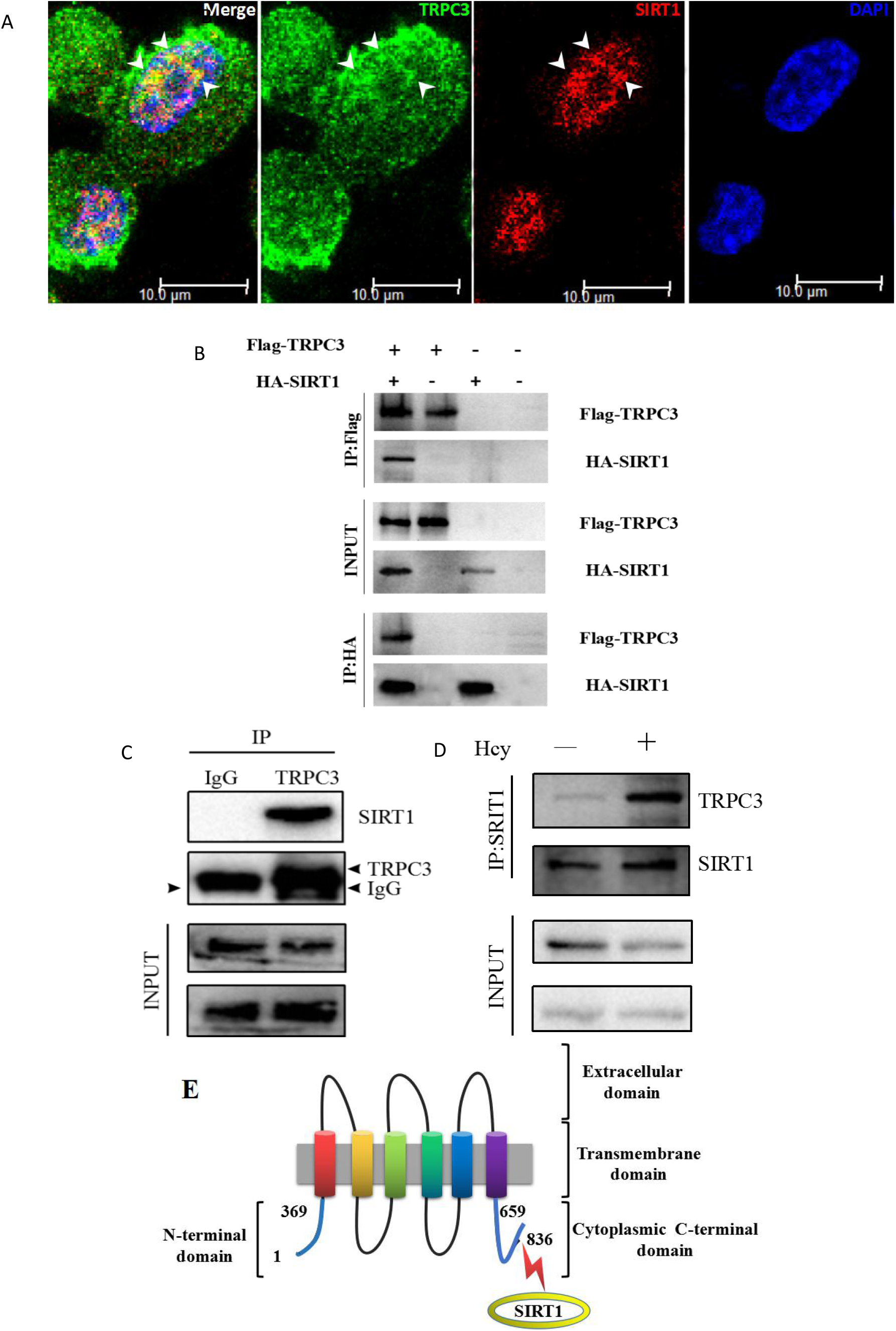
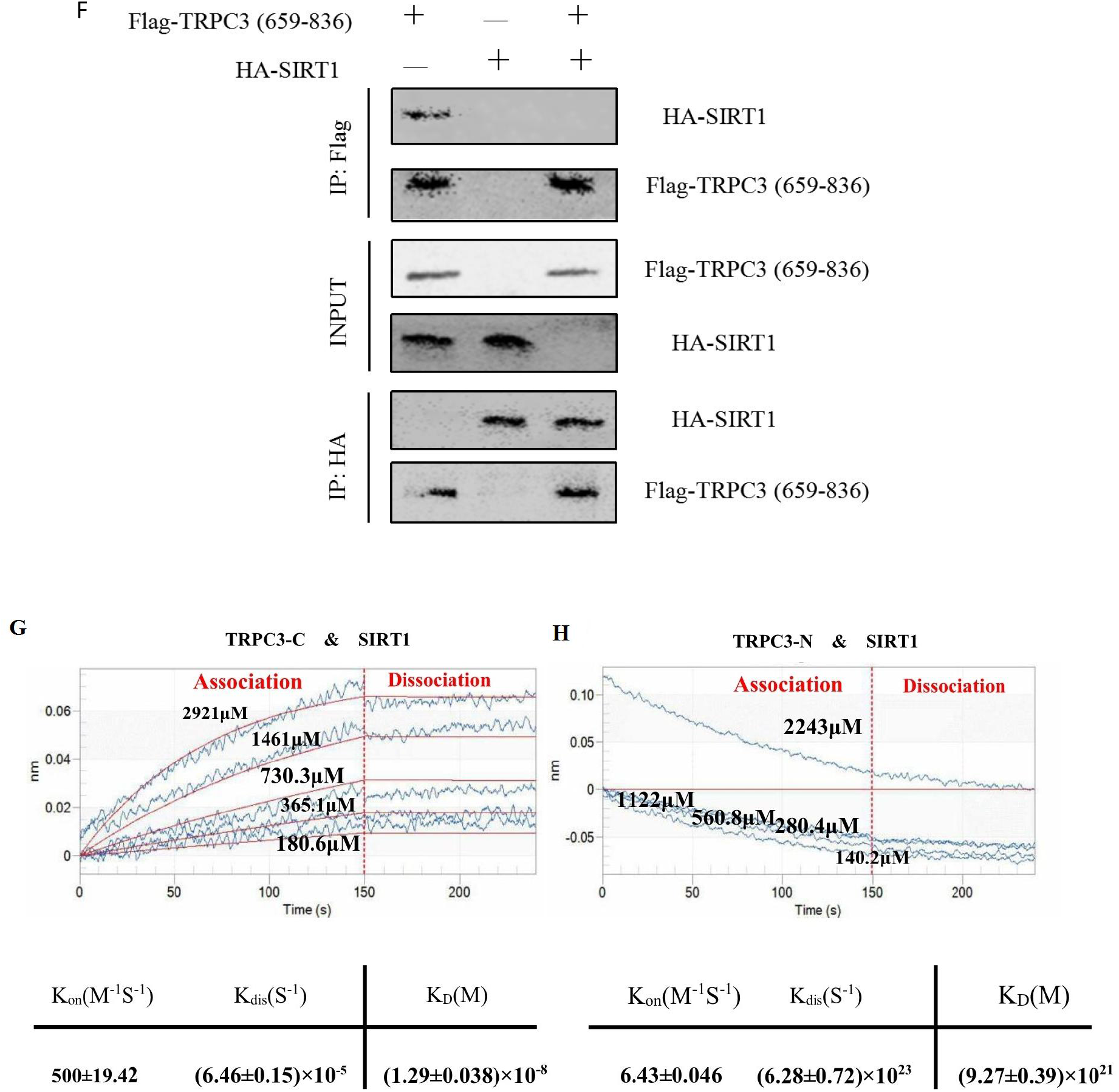
The interaction between TRPC3 and SIRT1. **A.** Co-localization of TRPC3 and SIRT1. Cells are co-stained with antibodies against TRPC3 (Alexa Fluor 488, green) and SIRT1 (Alexa Flour 564, red). Arrow indicates co-localization. **B.** Interaction of TRPC3 with SIRT1 in mammalian cells. **C.** Endogenous TRPC3 binds to SIRT1 in MCFs. **D.** The interaction of endogenous TRPC3 with SIRT1 in MCFs. **E.** Schematic illustration of the domains of TRPC3 and its association with SIRT1. **F.** The C-terminal tail (659-836) of TRPC3 interacts with SIRT1.**G.** and **H.** Analyse of the binding affinity between full-length SIRT1 and TRPC3-C (**G**) or TRPC3-N (**H**) by biolayer interferometry (BLI). All experiments were performed in triplicate.

Moreover, BLI results were used to evaluate the direct interaction between full-length SIRT1 and different binding fragments of TRPC3. TRPC3 contains three main domains, including extracellular, transmembrane and cytoplasmic N-terminal (1-369) or C-terminal (659-836) (Fig. 6E). The intensity of the binding affinity between purified protein of TRPC3-C and SIRT1 increased with various concentrations of TRPC3-C (180.6, 365.1, 730.3, 1461, 2921 μM) in a dose-dependent manner (Fig. 6G-6H). The values were presented as, K_on_ = 500 ± 19.42 M^−1^S^−1^ for the association phase, K_dis_= (6.46 ± 0.15) × 10^−5^ S^−1^ for the dissociation phase, and K_D_ = (1.29±0.038) × 10^−8^ M. However, the binding of SIRT1 and TRPC3-N was not existed, even if it had given different concentrations of TRPC3-N to response with SIRT1 peptide (140.2, 280.4, 560.8, 1122, 2243 μM). The curve was declined rather than ascended in the association. The values were K_on_ = 6.43 ±0.046 M^−1^S^−1^ for the association phase, K_dis_= (6.28 ± 0.72) × 10^23^ S^−1^ for the dissociation phase, and K_D_ = (9.27 ± 0.039) × 10^21^ M (Fig. 6H). In addition, This result was further verified in mammalian cells transfected with the plasmids encoding the C-tail of TRPC3 and full-length SIRT1 (Fig. 6F)

## 4. Discussion

With the increasing prevalence of cardiac structure remodeling in various cardiovascular diseases, the pathogenesis of cardiac fibrosisis an important research topic. Here we used hyperhomocysteinemia to aggravate the progression of atrial fibrosis and ultimately the structural re-entry circuits and local conduction block in the atrium.indirectly determine the basic pathogeny of AF. We found that patients with AF were more vulnerable to severe atrial fibrosis, than SR patients (Fig. 1E, Table1). Meanwhile, AF patients requiring cardiothoracic surgery were also more likely to develop hyperhomocysteinemia and HF as comorbidity (Table 1). TAC has proven to be a relatively reliable technique for generating HF models ^[24–26]^. In our study, TAC was performed at 4 weeks of age to establish HF models, which develop cardiac fibrosis; These mice were also fed a high-Hcy diet for 12-14 weeks to exacerbate the extent of cardiac fibrosis, which further demonstrated that Hcy triggers cardiac remodelling. As it is difficult to substantiate the spontaneous development of AF in mice, we applied acetylcholine in our experimental model ^[17]^, in order to observe the susceptibility of inducing arrhythmias by atrial fibrosis. This revealed that TAC and a high-Hcy diet increased the prevalence of AF compared to the NH mice, which we could confirm within 2 min of administering the injection of acetylcholine. A high Hcy diet alone may not cause cardiac structural changes in mice. For this reason, we use TAC to establish the model of HFpEF, which means that not only could we observe the relationship between cardiac fibrosis and a high Hcy diet in our HF model, but we could also detect the susceptibility of AF to Hcy stimulation in HF model. The increase in the incidence and duration of AF may be correlated with the extent of myocardial fibrosis in the TAC model ^[27,28]^. At the same time, there is likely a tendency to develop atrial fibrosis in the TAC animal models, moreover, a greater degree of fibrosis was observed in HH mice than in the NH group (Fig. 3I). This AF due to interruption of the continuity of atrial myocytes by excessive extracellular matrix (ECM) deposition and promotion of the formation of aberrant electrical circuits, which assisted in understanding the disorder of current conduction in AF (Fig. 3B). In our model, the duration of AF was more easily prolonged in the HH group than in the NH group, which further demonstrated that Hcy is a novel factor for atrial fibrosis-related AF and the increasing risk of AF ^[29,30]^. Among the treatment groups, HF mice were found tomanifest cachexia with different degree of death, which is characterized by a reduction in subcutaneous fat, activity, and food intake, starting from 7 weeks of age ^[31]^.

TRPC3 is considered an indispensable factor in regulating the mechanisms of fibrosis development in mouse cardiomyocytes. The expression level of this channel was higher in the left atrial appendage of AF patients than in SR patients, andaccompanied with enhance expression of the pro-fibrosis protein, TGF-ß. Concurrently, we founded similar results in the animal models; regardless of whether TAC mice were fed a high-Hcy diet or not, higher expression levels of TRPC3 and TGF-ß were observed in the HH mice compared with in the NH mice. Although the effect of added dietary folic acid and vitamins was not as notable as that achieved by directly knocking out the TRPC3 gene in HH mice, such dietary intervention may serve as a potential therapy for homocysteinemia patients with HF. Second, in our cell model, high-Hcy conditions elicited the up-regulation of TRPC3 protein and the pro-fibrosis modulator TGF-ß in a dose-dependent manner, thereby enhancing the proliferation and differentiation of fibroblasts.

We explored whether inhibition of TRPC3could attenuate the fibrotic response of cardiomyocytes induced by Hcy. Previous studies using TRPC3-deficient (TRPC3^−/−^) mice or a pharmacological inhibitor of TRPC3, Pyr-3, revealed that TRPC3 participated in mechanical stress-induced LV diastolic dysfunction in mice ^[32]^. Previous research has shown that TAC significantly increased myocardial cell size in both TRPC3^+/+^ and TRPC3^−/−^ mice. On the other hand, analysis of collagen deposition demonstrated a marked decrease in fibrosis in TRPC3^−/−^ mouse hearts compared to TRPC3^+/+^ mouse hearts ^[33]^. As interstitial fibrosis is perceived to be a major cause of cardiac remodelling, we assessed whether suppression of TRPC3 could attenuate Hcy-induced interstitial fibrosis in cardiac myocytes injured by TAC. The experimental results showed that the protein level of the fibrosis-related gene TGF-ß was significantly suppressed in TRPC3-KD mice after TAC (Fig. 4A-4D). Furthermore, in the cell models, We used MCF cell lines and atrial mouse fibroblasts respectively to confirm the diperacy between atrium and ventricle by the effect of Hcy.

The results showed that although the levels of fibrosis protein in both cells increased, the changes of atrial fibroblasts were more obvious. Next, we take atrial mouse fibroblasts as the cell model. Pyr-10, an inhibitor of TRPC3, could markedly control the extent of fibrosis in the cell models ^[33,34]^; TRPC3-shRNA-transfected fibroblasts exhibited attenuated Hcy-induced atrial fibrosis (Fig. 5B, 5D) and reduced the proliferation of fibroblasts (Fig. 5H). These results indicate that TRPC3 is critical for the proliferation and differentiation of fibroblasts and predominantly mediates Hcy-induced maladaptive fibrosis in mouse hearts. Generally, seven-transmembrane receptor TRPC3 is a critical mediator in the TGF-ß signalling pathway. Although TGF-ß signalling is crucial for cardiac remodelling ^[35]^, the role of TRPC3 in regulating TGF-ß and promoting fibrosis remains unknown.

We further studied the potential molecular mechanisms underlying how TRPC3 modulates TGF-ß-mediated cardiac fibrosis. It is worth acknowledging that up-regulation of SIRT1, an important cytokine that protects against cardiac remodelling, was observed and led to the inhibition of collagen accumulation by inhibiting TGF-ß expression ^[36,37]^. The severity of cardiac fibrosis was correlated with a lower expression level of SIRT1 in AF patients and TAC mice. However, adding Res, an activator of SIRT1, could also effectively attenuate cardiac fibrosis in both animal and cell models; strikingly, high doses of Res were also found to suppress TAC-induced fibrosis, especially obviously in the HH group (Fig. 4H). Similarly, SIRT1-overexpression mice efficiently mitigated the extent of atrial fibrosis compared with that in the SH+vector group (Fig. 4A-4D). Thus, SIRT1 may play a pivotal role in the mechanism of cardiac fibrosis.

Interestingly, SIRT1 overexpression inhibited the expression of TGF-ß, but no changes were not observed in the protein level of TRPC3 (Fig. 5E). In addition, with the increase of TRPC3 by titrating the amount of ectopic Flag-TRPC3, SIRT1 protein levels were significantly decreased in a dose-dependent manner, following the enhancement of TGF-ß (Fig. 5F), which suggested that SIRT1 might be required for the indirect adjustment of TGF-ß expression mediated by TRPC3. Indeed, application of the inhibitor of SIRT1, Salermide (Fig. 5B) to atrial fibroblasts further enhanced the protein level of TRPC3 and inhibition of SIRT1, as well as decreased TGF-ß expression under Hcy stimulation (Fig. 5B). Altogether, these results demonstrate that SIRT1 plays a crucial role in TRPC3-mediated TGF-ß signalling under Hcy stimulation and may play a role in protecting cell function and preventing cardiac remodelling.

The present study identified SIRT1 as a novel regulator and partner of TRPC3 via direct or indirect binding. Although previous studies have shown that both SIRT1 and TRPC3 participate in mediating TGF-ß signalling ^[32,38,39]^, it remains unknown how TPRC3 regulates the localization of SIRT1 and whether these proteins directly interact with each other. In fibroblasts transfected with purified TRPC3-shRNA lentiviruses, the expression of SIRT1 is increased with the MOI value in a dose-dependent manner, which indicates that the stability of SIRT1 is directly affected by TRPC3. Conversely, transfection of the SIRT1-overexpression vector into HEK293 cells did not affect the protein expression level of TRPC3 compared to transfection of a scrambled vector (Fig. 5K, 5L). Additionally, the co-IP, BLI and immunofluorescence results demonstrated the correlation between TRPC3 and SIRT1 in regulating the process of fibrosis (Fig. 6A-6D), and moreover, affirmed their binding domains to the C-terminal (659-836) of TRPC3 and SIRT1 (Fig. 6E-6H).

As further discussed below, TRPC3 can physically interact and subsequently activate SIRT1 in response to TGF-ß signalling and play a prominent role in promoting the proliferation and differentiation of fibroblasts. In this study, we investigated the mechanism of TRPC3 in the progression of atrial fibrosis both in vitro and in vivo. By analysing left atrial appendage specimens, we uncovered that TRPC3 levels were closely correlated with the degree of atrial fibrosis and the incidence of AF. Moreover, patients with hyperhomocysteinemia exhibited increased protein levels of TRPC3 and decreased levels of SIRT1 along with activation of the TGF-ß signalling pathway (Fig. 7). In agreement with this, our analysis showed that the RNA level of B was increased in the AF group, whereas the RNA level of A was reduced (Fig. 1C) compared with that in the SR group. Remarkably, we further found that the Hcy group exhibited a marked increase in fibroblast proliferation and enhanced expression of a pro-fibrosis protein, TGF-ß, in vitro and in vivo. However, even in the presence of different concentrations of Hcy, inhibition of TRPC3 led to a significant reduction in the proliferation and migration of fibroblasts (Fig. 5G, 5H). Interestingly, our experiments also revealed that the C-terminal domain of TRPC3 (residues 659-836aa) could act as a dominant-negative modulator to inhibit SIRT1.

**Figure 7.**
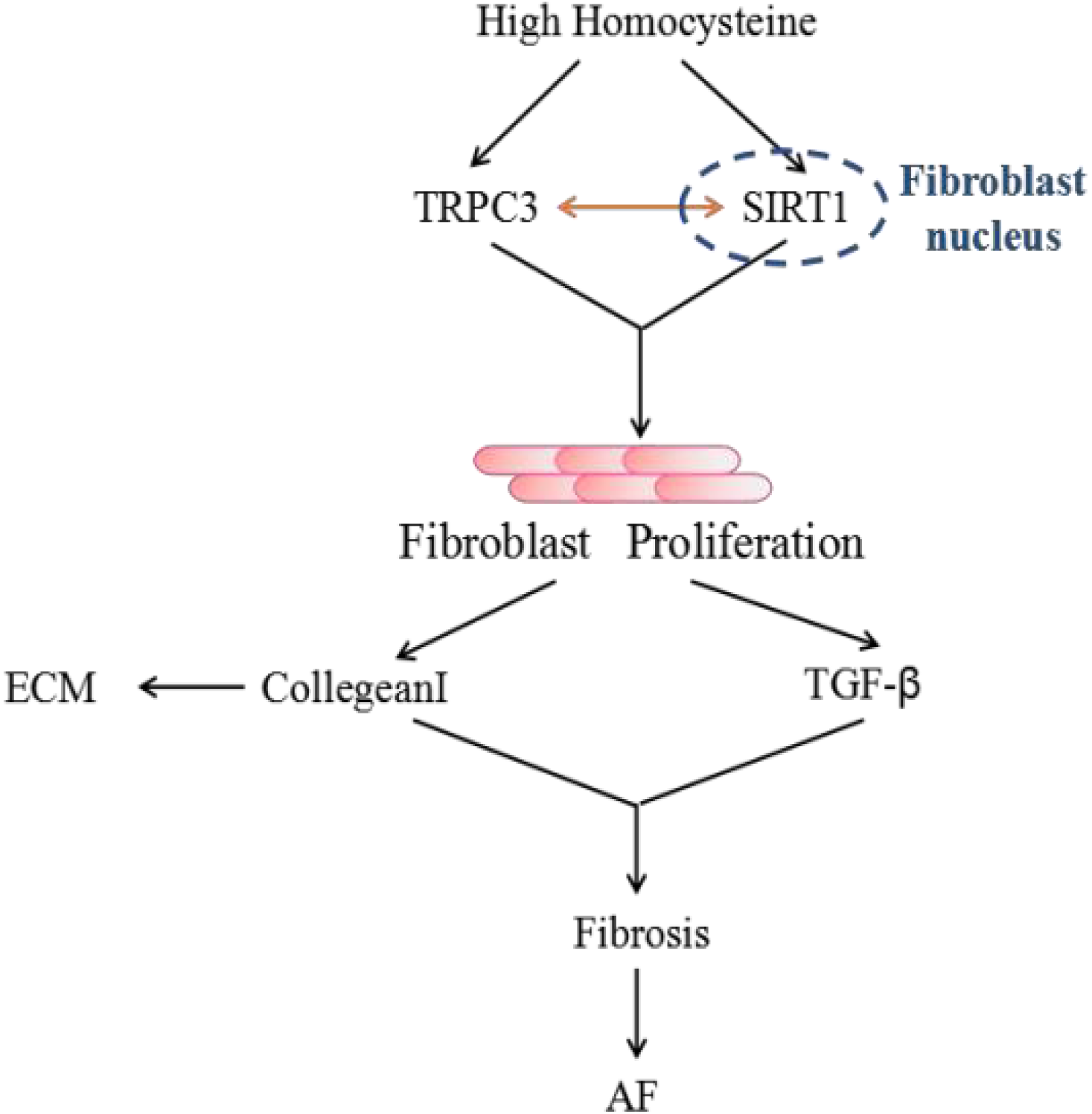
Schematic representation of our working hypothesis. Increased Hcy levels promote the proliferation and differentiation of fibroblasts, which can increase the protein levels of TPRC3 and through interaction with SIRT1 promote atrial fibrosis. This may occur though the direct activation of TGF-ß to induce collagen-I accumulation associated with the pathological mechanism of AF. However, the specific mechanism of AF requires further investigation.

## 5. Conclusions and Perspective

Our present study not only illustrates the biochemical function of TRPC3which binds directly to SIRT1 through its C-terminal (659-836aa) to modulate TGF-ß signaling, with the interaction intensified under Hcy stimulation, but also unveils the roles of TRPC3 and SIRT1 in atrial structure remodelling and fibrosis, which reversely increase the occurrence of AF accompanied by HF (Fig. 7). Finally, the changes among TRPC3 and TGF-ß are consistent on the transcript and protein levels and are in accordance with the changes in SIRT1 induced by Hcy. Together, our results suggest that TRPC3 may serve as a biomarker for preventing the consequence of atrial fibrosis in AF patients with homocysteinemia, especially as activation of SIRT1 is associated with inhibition of the TGF-ß signalling pathway. Thus, the underlying mechanisms of atrial fibrosis is mediated by TRPC3 and SIRT1 merits further investigation.

## Acknowledgements

We are grateful to Dr Juxiang Li for careful reading and revision of the manuscript.

---

**Supplement Figure 1 A.** Representative M-mode images of hearts of the treatment groups at 10 weeks of age. **B.** The ejection fraction (EF) level measured in the three groups. **C-D.** The Masson staining images (C) and Representative M-mode images (**D**) in different groups, SIRT1-overexpression(SIRT1-O)+SH, TRPC3-KD(TRPC3-D)+SH, SH and SH+vector, respectively.

**Supplement Figure 2 A.** MCF cells were infected with purified lentivirus for 24 hours, at multiplicity of infections (MOIs) of 2, 5, 10 and 20, respectively. **B.** Cell viability was not significantly affected by lentivirus transfection with or without Hcy stimulation

**Supplement Figure 3** Co-IP of purified protein of TRPC3-N, -C and SIRT1.

